# Fibrocyte accumulation in bronchi: a cellular hallmark of COPD

**DOI:** 10.1101/449843

**Authors:** Isabelle Dupin, Matthieu Thumerel, Elise Maurat, Florence Coste, Hugues Begueret, Thomas Trian, Michel Montaudon, Roger Marthan, Pierre-Olivier Girodet, Patrick Berger

## Abstract

**Background:** The remodeling mechanism and cellular players causing persistent airflow limitation in chronic obstructive pulmonary disease (COPD) remain largely elusive. We have recently demonstrated that circulating fibrocytes, a rare population of fibroblast-like cells produced by the bone marrow stroma, are increased in COPD patients during an exacerbation. It remains, however, unclear, whether fibrocytes are present in bronchial tissue of COPD patients.

**Objective:** We aimed to quantify fibrocytes density in bronchial specimens from both control subjects and COPD patients, and to define associations with clinical, functional and computed tomography relevant parameters.

**Methods:** 17 COPD patients and 25 control subjects with normal lung function testing and no chronic symptoms, all of them requiring thoracic surgery, were recruited. LFT and CT-scan were performed before surgery. Using co-immunostaining and image analysis, we identify CD45^+^ FSP1^+^ cells as tissue fibrocytes and quantify their density in distal and proximal bronchial specimens from the whole series.

**Results:** Here, we demonstrate that fibrocytes are increased in both distal and proximal tissue specimens of COPD patients, compared to those of controls. The density of fibrocytes is negatively correlated with lung function parameters, such as FEV1 and FEV1/FVC, and positively with bronchial wall thickness assessed by CT scan. High density of distal bronchial fibrocytes predicts presence of COPD with a sensitivity of 83% and a specificity of 70%.

**Conclusions:** Our results thus suggest that recruitment of fibrocytes in the bronchi may participate to lung function decline during COPD progression.

**Clinical Implications:** High density of tissue fibrocytes is associated with a deteriorated lung function and an increase in airway wall thickness. A low density tissue fibrocytes virtually eliminates the presence of COPD.

**Capsule summary:** Blood fibrocytes assessed during exacerbation is a predictor of mortality in COPD. This study shows an increase of bronchial fibrocytes, that is associated with lower lung function, increased bronchial thickness and air trapping in COPD.

## Introduction

Chronic obstructive pulmonary disease (COPD) is characterized by chronic persistent inflammation and remodeling leading to progressive airflow limitation (1, 2). The evolution of this chronic disease is worsened by acute exacerbations, frequently triggered by viral or bacterial infections (3). These exacerbations are considered as an independent prognostic factor for mortality (4). Current pharmacological treatments for COPD patients decrease exacerbation frequency by only up to 29% as compared to placebo either alone or in combination, but they do not have any significant effect on mortality (5–7). COPD patients exhibit remodeling processes leading to permanent changes in tissue structure, such as epithelial mucous metaplasia, parenchymal destruction (*i.e*., emphysema) and connective-tissue deposition in the small airway walls (1). This latter, also called peribronchiolar fibrosis, has been observed even in young smokers (8), thus suggesting that it may be an initiating event in COPD pathophysiology. To date, these processes are not inhibited or reversed by current pharmacotherapy.

Fibrocytes are fibroblast-like cells produced by the bone marrow stroma and released in the peripheral circulation (9). Circulating fibrocytes, defined as CD45^+^ collagen I ^+^ cells, are increased in COPD patients only during an exacerbation (10) and not in stable state, as compared to control subjects (10, 11). A high blood fibrocytes concentration during an exacerbation is associated with an increased risk of death (10), suggesting a deleterious role of fibrocytes in COPD evolution. By contrast, myeloid derived suppressor cells (MDSC)-like fibrocytes, a subpopulation of circulating fibrocytes, are increased in the blood of stable COPD patients, and these cells might rather play a protective role (11). The presence and the role of fibrocytes in the lung of COPD patients remain controversial (11) and need to be clarified. Indeed, tissue fibrocytes, defined as CD34^+^ collagen I^+^ cells, have not been found in distal airways of COPD patients, and detected in proximal airways of only less than 50% of COPD patients (11). However, fibrocytes are known to downregulate CD34 expression when differentiating (12). Thus, the co-expression of CD34 and collagen I, as a definition criterion for tissue fibrocyte, may lead to fibrocytes underestimation (11). As a consequence, we define fibrocytes as cells double positive for both CD45 and fibroblast-specific protein 1 (FSP1), in agreement with previous reports from human (13) and mice (14–16) lungs. Thus, the aim of the present study was to determine the density of tissue fibrocytes (*i.e*., CD45^+^ FSP1^+^ cells) in distal and proximal airway specimens of COPD patients, as compared to that in control subjects. We then evaluated the relationship between the density of tissue fibrocytes and parameters derived from lung function test and quantitative computed tomography (CT) as well as with blood fibrocytes. Functional *in vitro* experiments were also performed to assess the effect of epithelial microenvironment on fibrocyte survival.

## Methods

A more detailed description of methods is provided in the online supplement.

### Study populations

Subjects aged more than 40 years were eligible for enrolment if they required thoracic surgery lobectomy for cancer pN0, lung transplantation or lung volume reduction. A total of 17 COPD patients, with a clinical diagnosis of COPD according to the GOLD guidelines (2) and 25 non COPD subjects (“control subjects”) with normal lung function testing (*i.e*., FEV_1_/FVC > 0.70) and no chronic symptoms (cough or expectoration) were recruited from the University Hospital of Bordeaux.

To study fibrocytes *in vitro*, blood samples were obtained from a separate cohort of COPD patients, the COBRA cohort (“Cohorte Obstruction Bronchique et Asthme”; Bronchial Obstruction and Asthma Cohort; sponsored by the French National Institute of Health and Medical Research, INSERM) (Tables E3 and E5).

### Study design

This clinical trial was sponsored by the University Hospital of Bordeaux. The study has been registered at ClinicalTrials.gov under the N° NCT01692444 (*i.e.* “Fibrochir” study). The study protocol was approved by the local research ethics committee on May 30, 2012 and the French National Agency for Medicines and Health Products Safety on May 22, 2012. All subjects provided written informed consent. The study design is summarized in Fig E1. 4 visits were scheduled: a pre-inclusion visit (V1) to explain study and surgery, an inclusion visit (V2) the day of surgery, a visit one month ± 15 days after surgery (V3), and a the last visit one year ± 15 days after surgery (V4).

### Bronchial fibrocytes identification

A sub-segmental bronchus sample (for proximal tissue) as well as fragments of distal parenchyma were obtained from macroscopically normal lung resection material. The samples were embedded in paraffin and sections of 2.5 µm thick and were stained with both rabbit anti-FSP1 polyclonal antibody (Agilent) and mouse anti-CD45 monoclonal antibody (BD Biosciences, San Jose, CA), or mouse anti-CD3 monoclonal antibody (Agilent), mouse anti-CD19 monoclonal (Agilent), mouse anti-CD34 monoclonal antibody (Agilent). The sections were imaged using a slide scanner Nanozoomer 2.0HT (Hamamatsu Photonics, Massy, France). Quantification of dual positive cells for FSP1 and CD45 was performed as described in Fig 1. The density of FSP1^+^ CD45^+^ cells was defined by the ratio between the numbers of dual positive cells in the lamina propria divided by lamina propria area. Quantification of dual positive cells for FSP1 and CD3, FSP1 and CD19 or FSP1 and CD34 was performed, as described above with some modification (see Supplemental Material and Methods in the online data supplement). Tissue area and cell measurements were all performed in a blinded fashion for patients’ characteristics.

**Fig 1.**
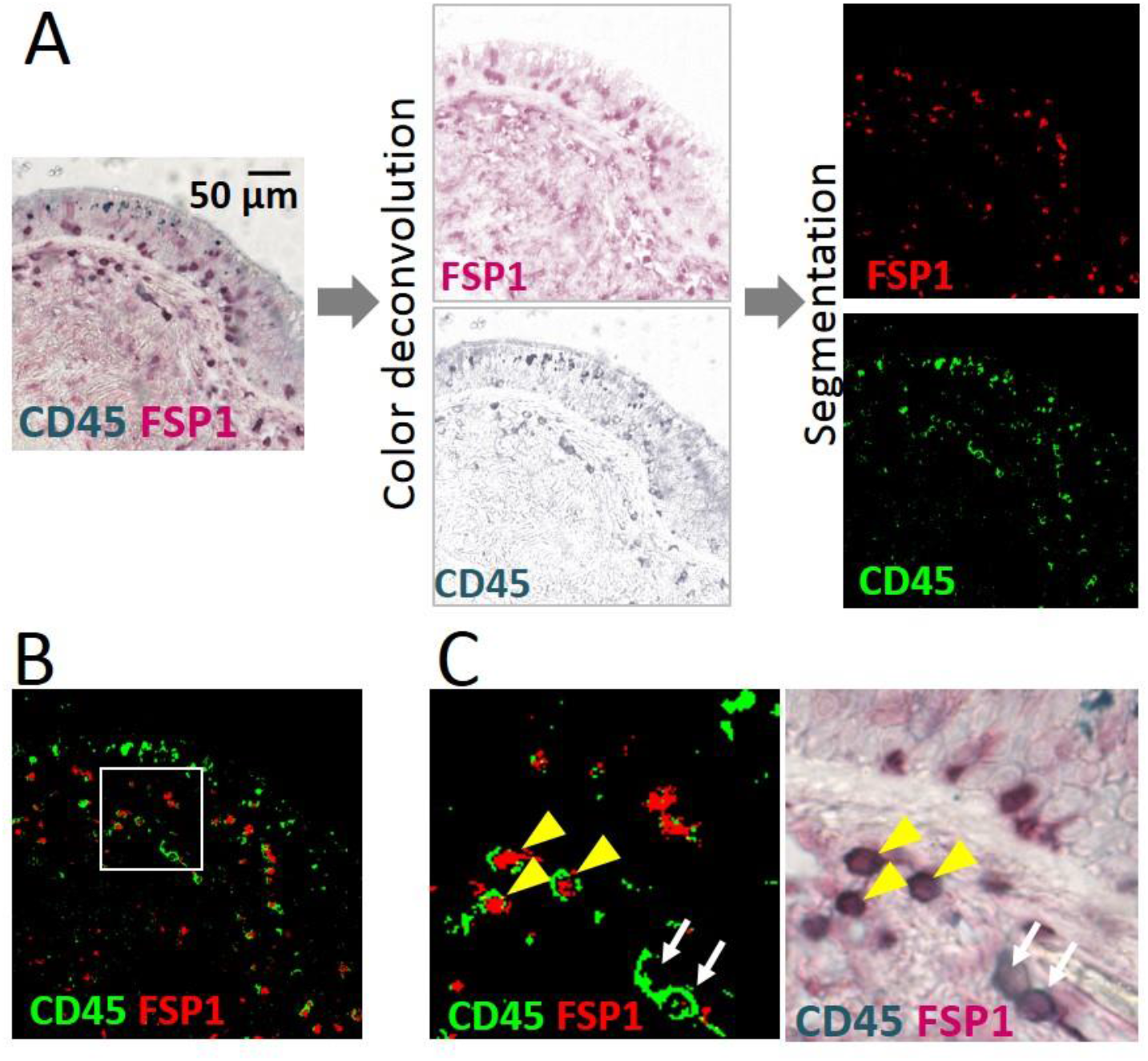
Detection of CD45^+^ FSP1^+^ cells. A, Left column, CD45 (green) and fibroblast-specific protein 1 (FSP1, red) stainings. Middle column, images for CD45 (top panel) and FSP1 (bottom panel) stainings are obtained after color deconvolution. Right column, segmented images are obtained for CD45 (top panel) and FSP1 stainings (bottom panel) after segmentation by a binary threshold. B, Merged segmented image. C, Higher magnification of the segmented image in B. The yellow arrowheads and the white arrows indicate respectively CD45^+^ FSP1^+^ cells and CD45^+^ cells.

### Quantitative computed tomography

CT scans were performed on a Somatom Sensation Definition 64 (Siemens, Erlangen, Germany) at full inspiration and expiration and analyzed using dedicated and validated software, as described previously (17–20)

### Circulating fibrocytes identification

Non-adherent non-T (NANT) cells were purified from Peripheral Blood Mononuclear Cells (PBMC) separated from the whole blood, and circulating fibrocytes were identified as double positive cells for the surface marker CD45 and the intracellular marker collagen I by flow cytometry, as described previously (10).

### Bronchial epithelial supernatants

Human bronchial epithelial cells (BEC) were derived from bronchial specimens (see Table E4 for patients’ characteristics), as described previously (21). Basal epithelial supernatant from fully diffenciated epithelium was collected for further experiments.

### Fibrocyte differentiation and survival

NANT cells purified from blood samples of COPD patients (Table E3 and E5) were incubated during one week in DMEM (Fisher Scientific) supplemented with 20% fetal calf serum (Biowest, Riverside, USA), followed by another week in serum-free medium, or serum-free medium containing 50% of basal epithelial supernatant. After 2 weeks in culture, the cells were detached 214 by accutase (Fisher Scientific), and either fixed overnight with Cytofix/Cytoperm and stained to 215 assess CD45, FSP1 and collagen I expression or directly stained by propidium iodide (PI) to assess 216 the level of dead cells (PI^+^ cells), by flow cytometry.

### Statistical analysis

Values are presented as means ± SD or the medians (95% confidence interval [CI]). Statistical 220 significance, defined as *P* < 0.05, was analyzed by Fisher’s exact tests for comparison of proportions, by two-sided independent t-tests for variables with a parametric distribution, and, by Wilcoxon tests, Mann-Whitney U tests and Spearman correlation coefficients for variables with a non-parametric distribution. Receiver operating characteristic (ROC) analysis and a univariate logistic regression analysis was performed to evaluate the association between COPD and a high density of tissue fibrocytes.

## Results

### Study population

The number of patients enrolled, excluded or followed for up to 1 year after surgery is shown in supplemental Fig E1. Clinical and functional characteristics, as well as quantitative CT parameters of all subjects with tissue fibrocytes assessment are shown in Table 1. The groups of control and COPD patients were well matched for age and body mass index. As expected, COPD patients were significantly different from controls in terms of smoking habits, lung function (FEV1, FVC, FEV1/FVC ratio, RV), diffusing capacity (TLCO) and CT parameters including wall thickness, emphysema extent (LAA), air trapping (MLA E) and cross-sectional pulmonary vessel area and number (CSA, CSN) (Table 1).

**Table 1:**
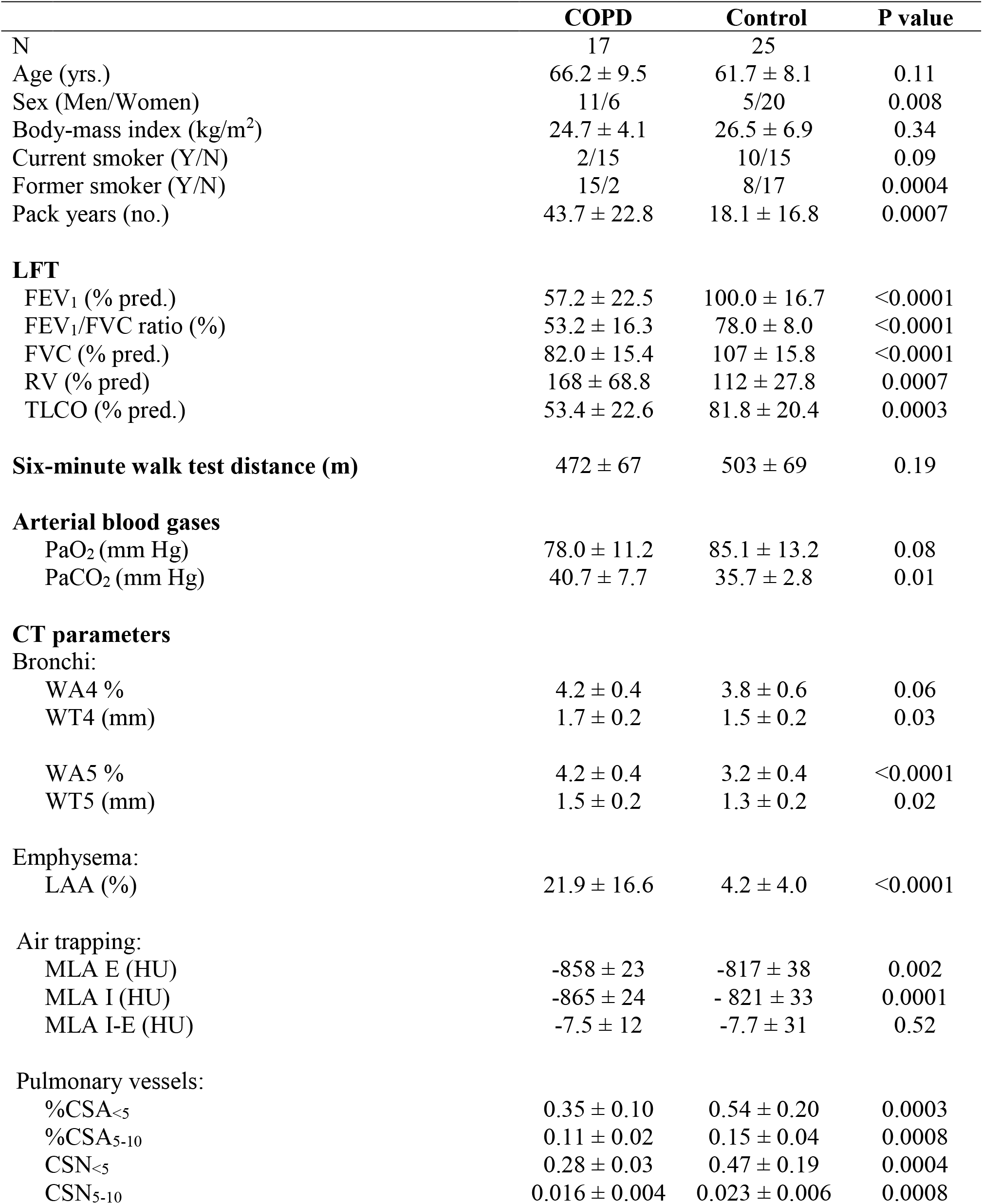
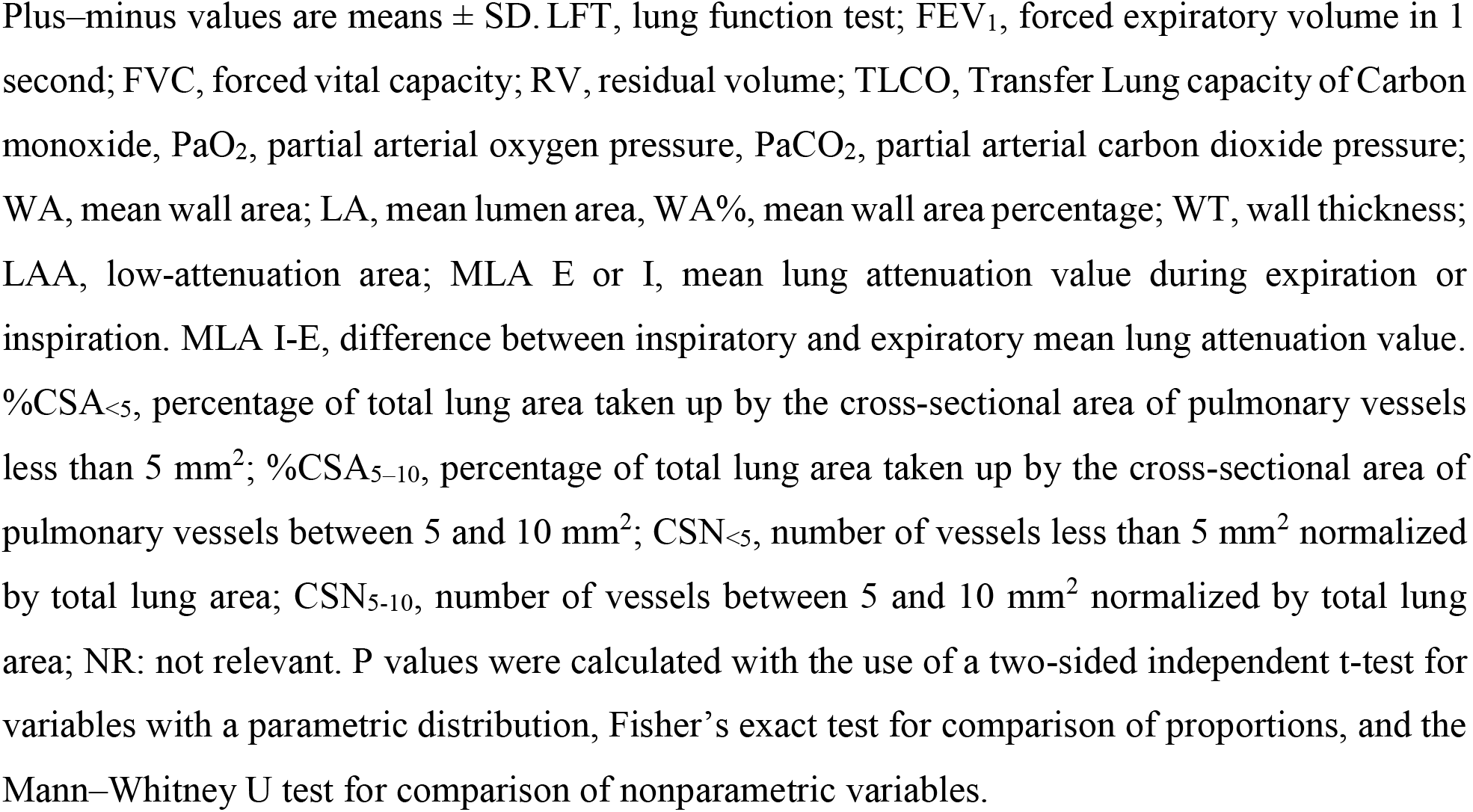
Patient characteristics

|  | **COPD** | **Control** | **P value** |
| --- | --- | --- | --- |
| **N** | 17 | 25 |  |
| Age (yrs.) | 66.2 ± 9.5 | 61.7 ± 8.1 | 0.11 |
| Sex (Men/Women) | 11/6 | 5/20 | 0.008 |
| Body-mass index (kg/m<sup>2</sup>) | 24.7 ± 4.1 | 26.5 ± 6.9 | 0.34 |
| Current smoker (Y/N) | 2/15 | 10/15 | 0.09 |
| Former smoker (Y/N) | 15/2 | 8/17 | 0.0004 |
| Pack years (no.) | 43.7 ± 22.8 | 18.1 ± 16.8 | 0.0007 |
| **LFT** |  |  |  |
| FEV<sub>1</sub> (% pred.) | 57.2 ± 22.5 | 100.0 ± 16.7 | <0.0001 |
| FEV<sub>1</sub>/FVC ratio (%) | 53.2 ± 16.3 | 78.0 ± 8.0 | <0.0001 |
| FVC (% pred.) | 82.0 ± 15.4 | 107 ± 15.8 | <0.0001 |
| RV (% pred) | 168 ± 68.8 | 112 ± 27.8 | 0.0007 |
| TLCO (% pred.) | 53.4 ± 22.6 | 81.8 ± 20.4 | 0.0003 |
| **Six-minute walk test distance (m)** | 472 ± 67 | 503 ± 69 | 0.19 |
| **Arterial blood gases** |  |  |  |
| PaO<sub>2</sub> (mm Hg) | 78.0 ± 11.2 | 85.1 ± 13.2 | 0.08 |
| PaCO<sub>2</sub> (mm Hg) | 40.7 ± 7.7 | 35.7 ± 2.8 | 0.01 |
| **CT parameters** |  |  |  |
| Bronchi: |  |  |  |
| WA4 % | 4.2 ± 0.4 | 3.8 ± 0.6 | 0.06 |
| WT4 (mm) | 1.7 ± 0.2 | 1.5 ± 0.2 | 0.03 |
| WA5 % | 4.2 ± 0.4 | 3.2 ± 0.4 | <0.0001 |
| WT5 (mm) | 1.5 ± 0.2 | 1.3 ± 0.2 | 0.02 |
| Emphysema: |  |  |  |
| LAA (%) | 21.9 ± 16.6 | 4.2 ± 4.0 | <0.0001 |
| Air trapping: |  |  |  |
| MLA E (HU) | -858 ± 23 | -817 ± 38 | 0.002 |
| MLA I (HU) | -865 ± 24 | -821 ± 33 | 0.0001 |
| MLA I-E (HU) | -7.5 ± 12 | -7.7 ± 31 | 0.52 |
| Pulmonary vessels: |  |  |  |
| %CSA<sub><5</sub> | 0.35 ± 0.10 | 0.54 ± 0.20 | 0.0003 |
| %CSA<sub>5-10</sub> | 0.11 ± 0.02 | 0.15 ± 0.04 | 0.0008 |
| CSN<sub><5</sub> | 0.28 ± 0.03 | 0.47 ± 0.19 | 0.0004 |
| CSN<sub>5-10</sub> | 0.016 ± 0.004 | 0.023 ± 0.006 | 0.0008 |

Plus-minus values are means ± SD. LFT, lung function test; FEV<sub>1</sub>, forced expiratory volume in 1 second; FVC, forced vital capacity; RV, residual volume; TLCO, Transfer Lung capacity of Carbon monoxide, PaO<sub>2</sub>, partial arterial oxygen pressure, PaCO<sub>2</sub>, partial arterial carbon dioxide pressure; WA, mean wall area; LA, mean lumen area, WA%, mean wall area percentage; WT, wall thickness; LAA, low-attenuation area; MLA E or I, mean lung attenuation value during expiration or inspiration. MLA I-E, difference between inspiratory and expiratory mean lung attenuation value. %CSA<sub><5</sub>, percentage of total lung area taken up by the cross-sectional area of pulmonary vessels less than 5 mm<sup>2</sup>; %CSA<sub>5-10</sub>, percentage of total lung area taken up by the cross-sectional area of pulmonary vessels between 5 and 10 mm<sup>2</sup>; CSN<sub><5</sub>, number of vessels less than 5 mm<sup>2</sup> normalized by total lung area; CSN<sub>5-10</sub>, number of vessels between 5 and 10 mm<sup>2</sup> normalized by total lung area; NR: not relevant. P values were calculated with the use of a two-sided independent t-test for variables with a parametric distribution, Fisher's exact test for comparison of proportions, and the Mann-Whitney U test for comparison of nonparametric variables.

### Bronchial fibrocytes are increased in COPD patients

As a methodological control, we first cultured fibrocytes from blood samples coming from a separate cohort of COPD patients (Table E3), and we showed that virtually all the CD45^+^ FSP1^+^ cells (99.9 ± 0.06%) purified from circulating PBMC also express collagen I after 14 days of cell differentiation *in vitro* (Fig E2). This allows us to define tissue fibrocytes as CD45^+^ FSP1^+^ cells. These cells were identified by immunohistochemistry as shown in Fig 1, and they were detected in distal tissue specimens from 11 of 12 COPD patients (92%) and 13 of 20 control subjects (65%) (Fig 2) as well as in proximal tissue specimens from 14 of 14 COPD patients (100%) and 16 of 21 control subjects (76%) (Fig 3).

**Fig 2.**
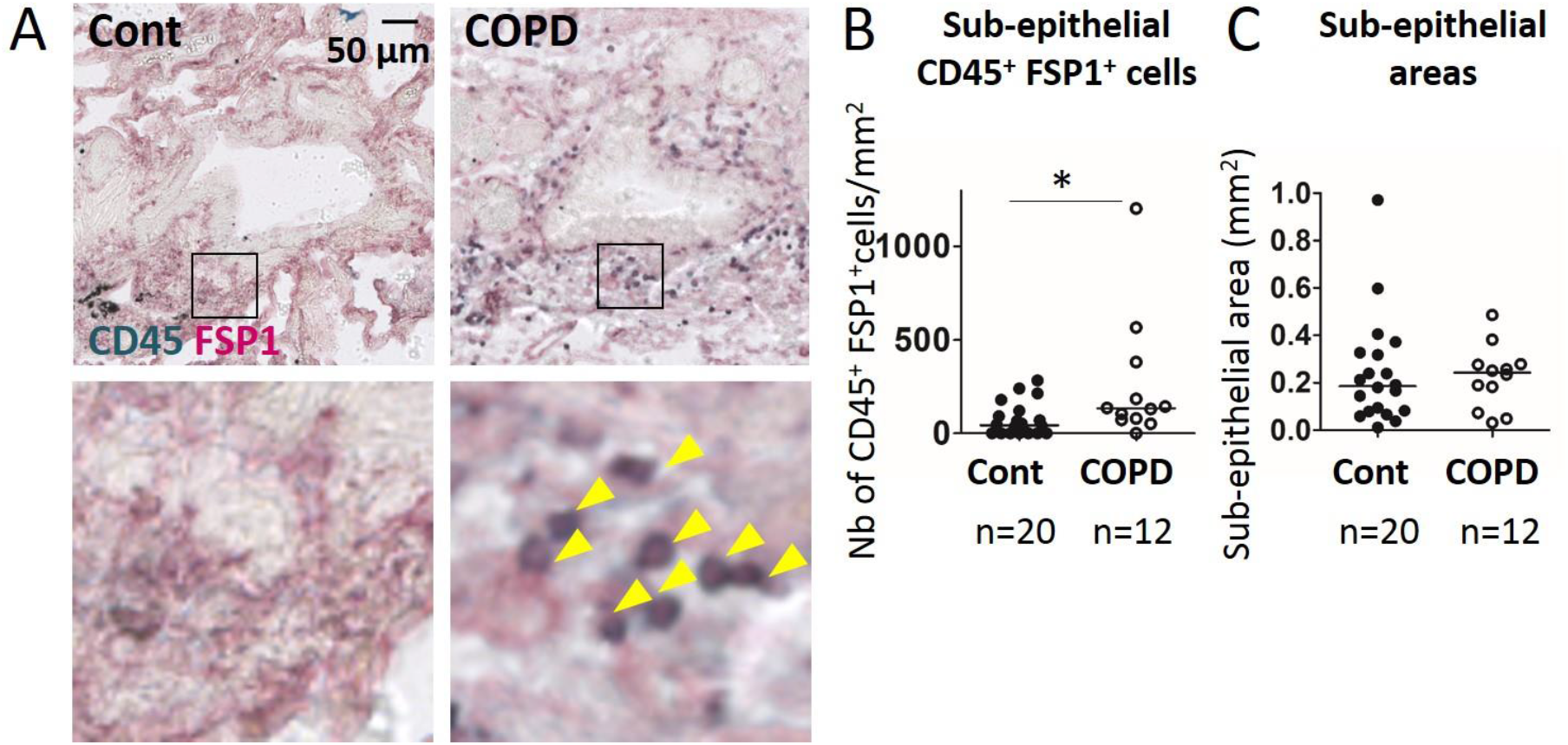
Increased fibrocyte density of in distal airways of COPD patients. A, Representative staining of CD45 (green) and FSP1 (red) in distal bronchial tissue specimens from control subject (left) and COPD patient (right). The yellow arrowheads indicate fibrocytes, defined as CD45^+^ FSP1^+^ cells. B, Quantification of fibrocyte density (normalized by the sub-epithelial area) in one specimen/patient. *: P<0.05, Mann Whitney test. C, Comparison of sub-epithelial areas in control subjects and COPD patients. B, C, medians are represented as horizontal lines.

These fibrocytes were located in the sub-epithelial region of both distal and proximal airways (Figs 2A and 3A) and, occasionally, within the epithelial layer. No CD45^+^ FSP1^+^ cell was evidenced within the airway smooth muscle layer. Some tissue fibrocytes were found in peribronchial area outside the smooth muscle layer (Fig E3). However, the analysis of fibrocyte density in this latter region could not be performed systematically, since this area could not be identified in each of our tissue specimens. The density of bronchial fibrocytes was higher in the subepithelial region of distal airways from COPD patients (median = 133 cells/µm^2^ (95% CI, 40 to 469), n = 12) than in that of control subjects (median = 42 cells/µm^2^ (95% CI, 31 to 114), n = 20, P<0.05) (Fig 2B). Similarly, fibrocytes density was also increased in the proximal airways from COPD patients (median = 73 cells/µm^2^ (95% CI, 47 to 139), n = 14) compared with control subjects (median = 21 cells/µm^2^ (95% CI, 18 to 60), n = 21, P<0.05) (Fig 3B). In both distal and proximal airways, there was no difference in sub-epithelial areas considered for tissue fibrocytes quantification between COPD patients and control subjects (Figs 2C and 3C). Not surprisingly, however, the density of fibrocytes in the sub-epithelial area of proximal tissue was positively and significantly correlated with that measured in distal airways (Fig E4).

**Fig 3.**
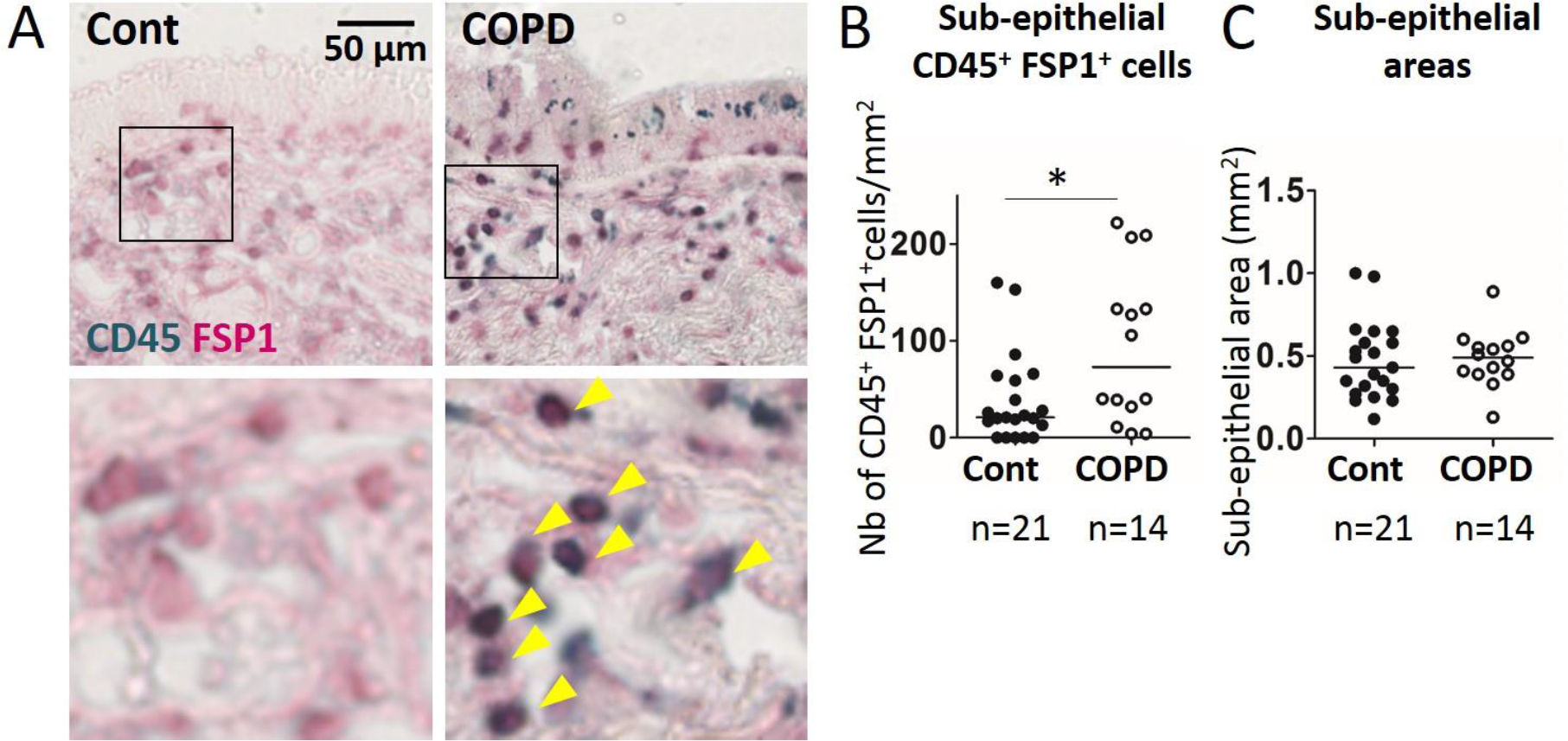
Increased fibrocyte density in proximal airways of COPD patients. A, Representative staining of CD45 (green) and FSP1 (red) in proximal bronchial tissue specimens from control subject (left) and COPD patient (right). The yellow arrowheads indicate fibrocytes, defined as CD45^+^ FSP1^+^ cells. B, Quantification of fibrocyte density (normalized by the sub-epithelial area) in one specimen/patient. *: P<0.05, Mann Whitney test. C, Comparison of sub-epithelial areas in control subjects and COPD patients. B, C, medians are represented as horizontal lines.

To further confirm our results, we co-stained FSP1 with CD3 or CD19 to determine whether FSP1 positive cells could be T-lymphocytes or B-lymphocytes, respectively. Except for one control subject, very few CD3^+^ cells also expressed FSP1 (Fig E5A), and there was no significant difference in the density of CD3^+^ FSP1^+^ cells in the sub-epithelial region of distal airways between controls and COPD patients (Fig E5B-C). Likewise, this density was not statistically different between both groups in proximal airways (Fig E5D-E). The B-lymphocyte CD19 marker co-localized with FSP1 positive cells neither in distal (Fig E6A) nor in proximal (Fig E6B) tissue specimens. We also co-immunostained CD34 and FSP1. CD34^+^ FSP1^+^ cells were detected in distal tissue specimens, from only 2 of 12 COPD patients (17%) and 4 of 20 control subjects (20%) (Fig E7). CD34^+^ FSP1^+^ cells were found in proximal tissue specimens from 9 of 13 COPD patients (69%) and 11 of 21 control subjects (52%) (Fig E8), but the density of these cells in COPD patients (median = 0.5 cells/µm^2^ (95% CI, 0.1 to 1.7), n = 13) was very low compared with that of CD45^+^ FSP1^+^ cells (median = 73 cells/µm^2^ (95% CI, 47 to 139), n=14) (Fig 2B). In both distal and proximal airways, there was no difference in the density of CD34^+^ FSP1^+^ cells between COPD patients and control subjects (Figs E7 and E8).

### Relationships between bronchial fibrocytes density and functional and CT parameters

We first determined univariate correlation coefficients between the density of tissue fibrocytes in the subepithelial region of both distal and proximal airways and various functional and CT parameters (Tables E1 and E2). In distal tissue specimens, the density of fibrocytes was negatively correlated to FEV1/FVC ratio (Fig 4A) and positively to PaCO2 (Fig 4B). It was also significantly associated with mean lung attenuation value during exhalation (Fig 4C). In proximal tissue specimens, the density of fibrocytes was negatively correlated to FEV1 (Fig 4D), FVC (Table E2), and positively correlated to RV (Table E2), WT4 (Fig 4E), WT5 (Fig 4F), WA4% (Table E2). Receiver Operator Characteristic (ROC) curves were built for all subjects whose density of tissue fibrocytes has been assessed in distal (n = 12 COPD patients and 20 control subjects, Fig 5A) and proximal (n = 14 COPD patients and 21 control subjects, Fig 5B) tissue specimens with significant areas under the curves (Table 2). To predict COPD, the density of fibrocytes in distal airways has a sensitivity of 83% and a specificity of 70%, whereas this density has a sensitivity of 79% and a specificity of 67% in proximal airways (Table 2). Moreover, the negative predictive value to eliminate COPD was 97.5% and 96.6% for distal and proximal airways, respectively, using a prevalence of 10% for COPD in the general population (22). ROC analyses allowed us to select the optimal value of fibrocytes density (cut-off values of 72 and 32 for distal and proximal tissue, respectively) to classify patients either with a high or low level of tissue fibrocytes (Table 2). COPD was associated with a high density of fibrocytes in distal (odds ratio: 11.7; 95% CI: [1.9-70.2]; P < 0.05) and proximal (odds ratio: 7.3; 95% CI: [1.5-35.1]; P < 0.05) airways. Thus, a high tissue fibrocytes density is associated with a higher risk of COPD.

**Fig 4.**
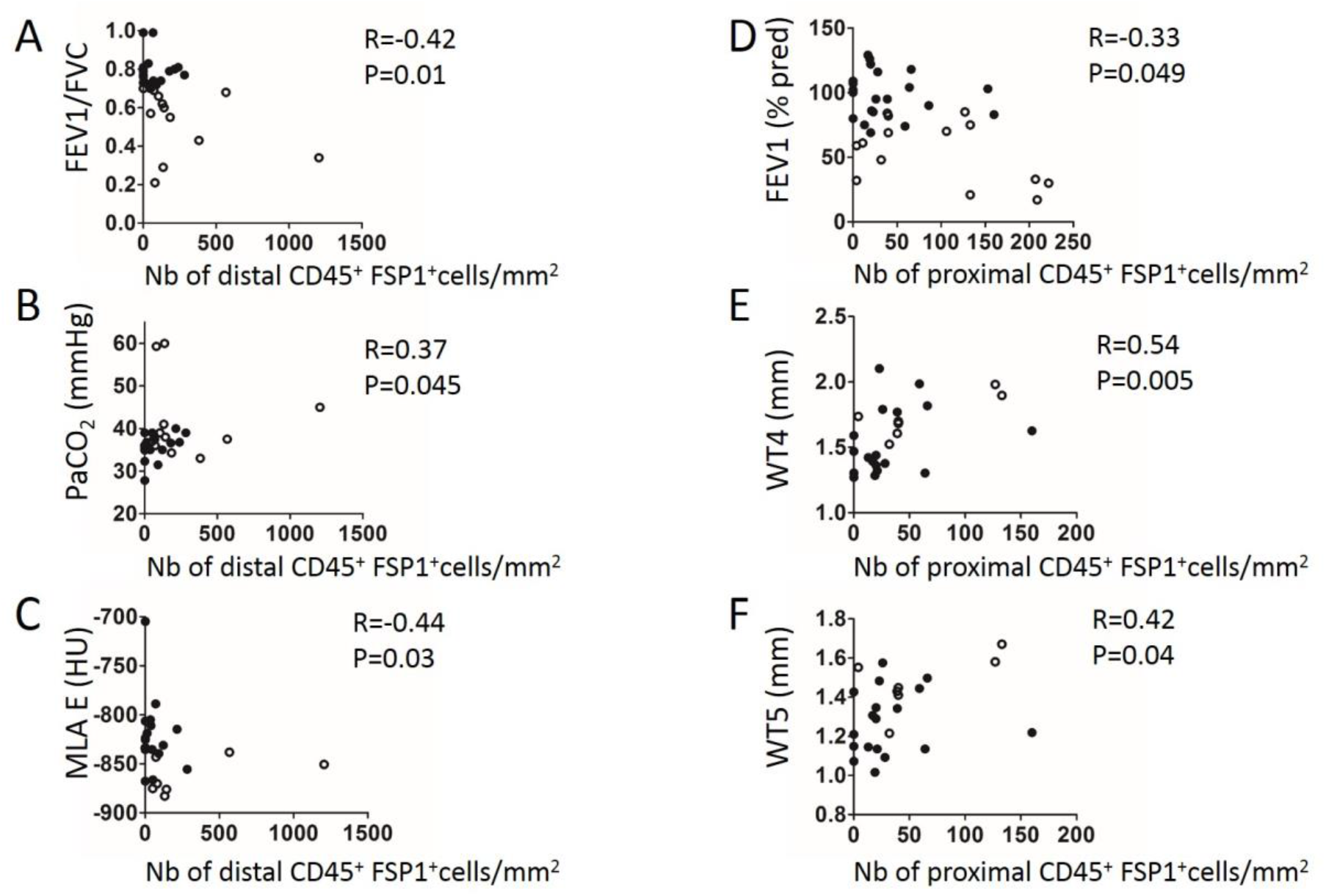
Relationships between fibrocyte density, lung function parameters and CT parameters. A-C, Relationships between FEV1/FVC (A), PaO_2_ (B), MLA during expiration (C) and the density of CD45+ FSP1+ cells in distal airways measured in control subjects (black circles) and COPD patients (open circles). D-F, Relationships between FEV1 (D), WT4 (E), WT5 (F) and the density of CD45+ FSP1+ cells in proximal airways measured in control subjects (black circles) and COPD patients (open circles). Correlation coefficient (r) and significance level (P value) were obtained by using nonparametric Spearman analysis.

**Fig 5.**
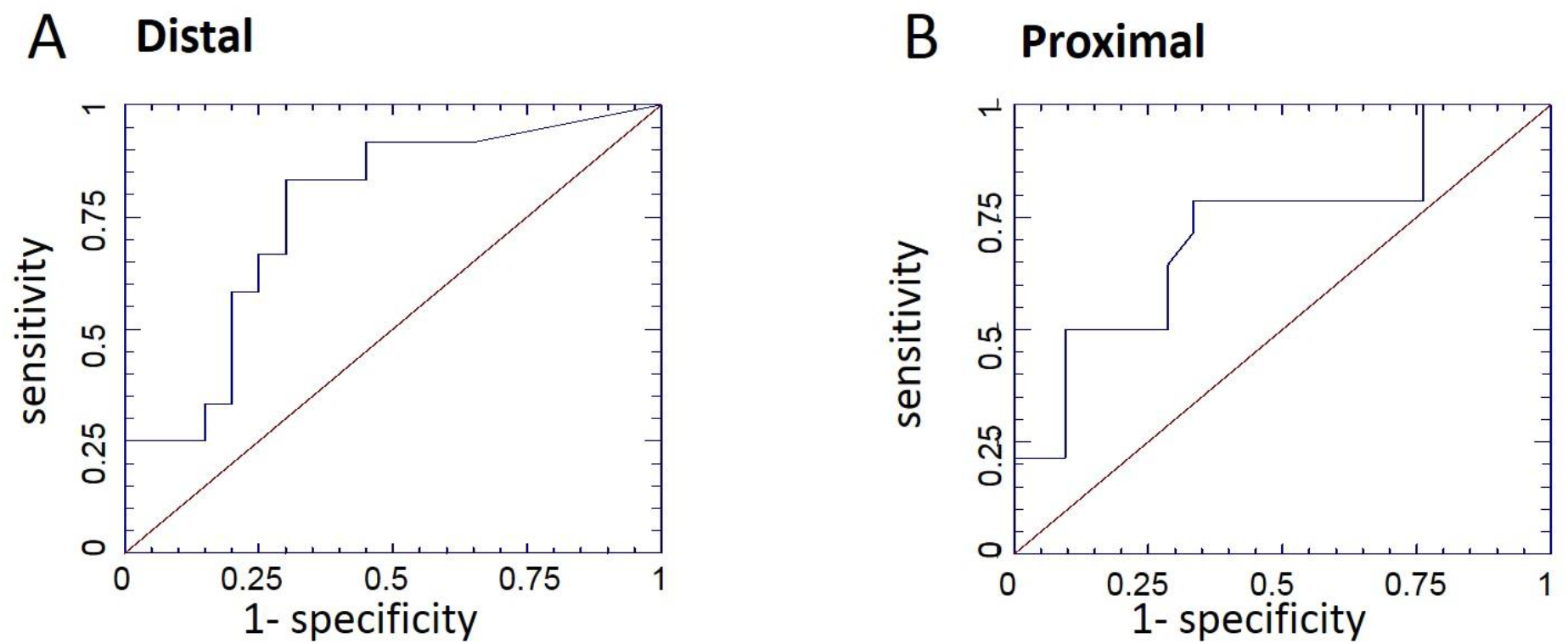
Diagnosis accuracy of high fibrocyte density for COPD. A-B, Receiver operating characteristic (ROC) curves for control subjects and COPD patients with 549 fibrocyte density measured in distal (A) or proximal (B) tissue specimens was built in order to 550 predict COPD.

**Table 2:** Association of high level of density of tissue fibrocytes with COPD

| Type of tissue | n | AUC ± SD | P value | Cut-off value | Sensitivity [IC] | Specificity [IC] |
| --- | --- | --- | --- | --- | --- | --- |
| **Distal** | 32 | 0.76 ± 0.09 | 0.04 | 72 | 0.83<br>[0.70-0.96] | 0.70<br>[0.54-0.86] |
| **Proximal** | 35 | 0.72 ± 0.09 | 0.02 | 32 | 0.79<br>[0.65-0.92] | 0.67<br>[0.51-0.82] |

Data are absolute number with [95% confidence interval] and plus–minus values are means ± SD. AUC, Area Under ROC Curve; ROC, Receiver Operating Characteristic; SD, standard deviation. P values, sensitivity and specificity were evaluated for ROC curve analysis.

### Circulating fibrocytes are unchanged in stable COPD

The percentage of blood fibrocytes (CD45^+^ ColI^+^ cells) in PBMC, was not statistically different in stable COPD patients (median=10.3% (95% CI, 4.6 to 16.5) of PBMC, n=12) and control subjects (median=7.9% (95% CI, 4.1 to 11.6) of PBMC, n=22) (Fig E9A). A similar result was obtained when fibrocytes concentration is expressed as absolute counts per milliliter of blood (data not shown). Finally, the percentage of blood fibrocytes (*i.e*., CD45^+^ ColI^+^ cells) in PBMC was significantly correlated with the density of bronchial fibrocytes (*i.e*., CD45^+^ FSP1^+^ cells) in distal airways (Fig E9B).

### COPD epithelial supernatant favors fibrocytes survival

We next investigated whether secretion from bronchial epithelial cells (BEC) from control or COPD patients could affect fibrocytes viability or ECM secretion in an *in vitro* assay. We evaluated this effect using BEC obtained from lung resection material sampled either in control subjects (n=2) or in COPD patients (n=2) (Table E4), cultured at the air-liquid interface. Fibrocytes were cultured from blood samples coming from a separate cohort of 6 COPD patients (Table E5). 7 to 10 days after blood sampling, cells, almost all being CD45^+^ FSP1^+^ cells (94.9 ± 3.6%), were exposed during 7 days to a mixture of fully differentiated BEC supernatants coming either from control subjects or COPD patients. 7 days after initial exposure, the level of CD45^+^ FSP1^+^ cells remains high (92.9 ± 3.9% and 93.4 ± 3.3% respectively for the control and COPD conditions). However, exposure of fibrocytes to COPD epithelial supernatant significantly decreased the percentage of dying cells (Fig E10).

## Discussion

In the present study, we have shown that the density of tissue fibrocytes (*i.e*., CD45^+^ FSP1^+^ cells) is significantly greater in both distal and proximal airway specimens of COPD patients, as compared to that of control subjects. We also found a significant correlation between this tissue fibrocytes density with blood fibrocytes as well as airflow obstruction, increased wall thickness, or air trapping. By means of ROC curve analysis and univariate logistic regression analysis, we observed that a high density of tissue fibrocyte increases the likelihood of COPD. Finally, it appears that fibrocytes survival is increased by epithelial cells secretion from COPD patients, a mechanism that could contribute to the elevated sub-epithelial density of fibrocytes in COPD patients.

There is an discrepancy between the present results and those previously obtained by Wright and colleagues (11), which deserves some methodological discussion. Wright *et al* did not find an increased level of tissue fibrocytes in COPD patients. They did not even observe any fibrocyte in distal airways (11). However, the methodology of tissue fibrocytes assessment and, ultimately, the definition of fibrocytes, are different in the present study and in that of Wright *et al* (11). Wright and colleagues’ method relied on the identification of CD34 and collagen I staining on sequential, instead of identical, sections (11) which could lead to either, false fibrocytes identification because of apparent co-expression in closely apposed but not identical cells, or fibrocytes underestimation because of the absence of co-expression in cells that are present only in one section. We have carefully addressed this issue by using antibodies and chromogens that are compatible with co-immunostaining in the same section. We have thus developed an image analysis technique that unambiguously identifies fibrocytes. Moreover, defining tissue fibrocytes as CD34^+^ collagen I^+^ cells (11) may lead to fibrocytes underestimation, which is consistent with previous data showing a downregulation of CD34 expression when fibrocytes differentiate in culture (12). In the connection, the present data confirming the very low density of CD34^+^ FSP1^+^ cells in bronchial specimens thus are in agreement with those of Wright *et al*. Fibrocytes are commonly defined as cells co-expressing CD45 and collagen I (23). The expression of a hematopoietic marker, such as CD45, is one of the minimum criteria for fibrocyte identification (23) and has been used in the present study. However, since immunohistochemistry for collagen I fails to unambiguously identify collagen I ^+^ cells in our experiments (data not shown), as well as in another report (24), we rather used the FSP1 marker, as a co-marker with CD45 for fibrocyte identification, as previously extensively described (13–16). FSP1 (25), also known as S100A4, is used as a marker for lung fibroblasts (24). Most of pulmonary FSP1^+^ cells express collagen I (24, 26), and the number of FSP1^+^ cells correlates with the extent of lung fibrosis in a murine model of fibrosis, suggesting that these cells contribute to collagen deposition (24). In addition, we have shown that almost all the CD45^+^ FSP1^+^ cells purified from circulating PBMC also express collagen I after 14 days of cell differentiation *in vitro* (Fig E2). It thus appears that the term of tissue fibrocyte is more accurate for double positive cells for CD45 and FSP1. Since FSP1 expression has been initially identified in fibroblasts (25) but was subsequently characterized in immune cells such as T and B lymphocytes (13), we paid a special attention to co-immunostain FSP1 with CD3, or CD19. In doing this, we showed that T lymphocytes co-expressing CD3 and FSP1 represented only a minor subset of CD45^+^ FSP1^+^ cells, the density of which did not change in COPD patients. We also showed that no B lymphocyte co-expressing CD19 and FSP1 was present in either distal or proximal human bronchi from both control subjects and COPD patients. Finally, FSP1 expression has also been characterized in macrophages (26). However, numerous macrophage markers, such as CD68, CD163, CD204, CD206, CD209 are also expressed by fibrocytes (27–29). It is thus impossible to properly differentiate fibrocytes from macrophages using immunohistochemistry even if a collagen I^+^ CD45^+^ double staining would have been suitable.

Our results may argue in favor of a potential deleterious role of tissue fibrocytes in COPD. Indeed, the greater the density of bronchial fibrocytes, the lower the FEV1 value or the FEV1/FVC ratio. Likewise, the greater the density of bronchial fibrocytes, the larger the bronchial wall thickness or the pulmonary air trapping. We previously pointed out a potential detrimental role of blood fibrocytes, since a high concentration of fibrocytes, in the peripheral circulation of COPD patients during an acute exacerbation, was associated with a higher mortality (10). Moreover, blood fibrocytes were present at a high level in frequent exacerbating patients (10). Since small airways are the major site of airway obstruction in COPD (30–32), the observation that tissue fibrocytes density, assessed in the present study, was higher in distal than in proximal airways makes sense. The correlation between the concentration of circulating fibrocytes and the density of distal bronchial fibrocytes may indicate that fibrocytes, which are recruited in the blood during an acute exacerbation (10), subsequently migrate to the airways and participate to the tissue remodeling process, such as peribronchial fibrosis leading to airway obstruction and air trapping. Indeed, once recruited into the lungs, fibrocytes may play various roles, including matrix secretion and degradation, pro-fibrotic cytokines production and activation of contractile force (33). Finally, the lower number of cell death in fibrocytes cultured with BEC from COPD patients combined with the elevated density of fibrocytes at the proximity of the epithelium in tissue samples from COPD patients, would suggest that secretion from epithelial cells in a COPD microenvironment provide pro-survival signals for tissue fibrocytes and could explain, at least partially, bronchial accumulation of fibrocytes in COPD patients. However, continuous efforts are warranted to clarify the cellular mechanisms by which fibrocytes may participate to obstruction development.

The present study has some limitations, which deserve further comments. The specimens for fibrocytes detection were obtained from patients with a diagnosis of lung cancer in 37 of 42 patients. As FSP1 is often expressed in malignant cells (34, 35), one may suggest that it could have an impact on fibrocytes density measured in our study. Since (i) bronchial specimens for fibrocytes analysis were selected from macroscopically normal lung resection material, and (ii) patients with a staging different from pN0 confirmed after surgery were excluded, it is unlikely that malignancy would have been sufficient to explain the increase in fibrocytes density we observed in COPD patients.

In conclusion, by taking advantage of unambiguous fibrocytes identification by co-immunostaining and image analysis, we unveil that COPD patients exhibit a greater density of fibrocytes in distal and proximal airways than control subjects. A high density of tissue fibrocytes is associated with a degraded lung function, airway structural changes and a higher risk of COPD, suggesting a deleterious pathogenic role for fibrocytes in COPD.

## Acknowledgments

The authors thank the study participants, the staff of thoracic surgery, radiology, pathology, respiratory and lung function testing departments from the University Hospital of Bordeaux, Virginie Niel for technical assistance, the Bordeaux Imaging Center (BIC) for help with imaging and image analysis. Microscopy was done in the Bordeaux Imaging Center a service unit of the CNRS-INSERM and Bordeaux University, member of the national infrastructure France BioImaging supported by the French National Research Agency (ANR-10-INBS-04). The help of Christel Poujol, Sèbastien Marais and Fabrice Cordelières for imaging and the help of Muriel Cario-Andrè for immunohistochemistry are acknowledged.

## Online Data Supplement

### Supplemental Material and Methods

#### Study Populations

Subjects aged more than 40 years were eligible for enrolment if they required thoracic surgery for lobectomy for cancer pN0, lung transplantation or lung volume reduction. A total of 17 COPD patients, with a clinical diagnosis of COPD according to the GOLD guidelines (1) and 25 non COPD subjects (“control subjects”) with normal lung function testing (*i.e*., FEV1/ FVC > 0.70) and no chronic symptoms (cough or expectoration) were recruited from the University Hospital of Bordeaux. Main exclusion criteria for both COPD patients and control subjects were history of asthma, lung fibrosis, idiopathic pulmonary hypertension and chronic viral infections (hepatitis, HIV). The main withdrawal criterion for subjects included for lobectomy due to cancer was a staging different from pN0 confirmed after surgery.

To study fibrocyte survival *in vitro*, blood samples were obtained from a separate cohort of COPD patients. These patients were recruited from the COBRA cohort (“Cohorte Obstruction Bronchique et Asthme”; Bronchial Obstruction and Asthma Cohort; sponsored by the French National Institute of Health and Medical Research, INSERM), as outpatients in the Clinical Investigation Centre of the University Hospital of Bordeaux (Tables E3 and E5).

To assess the role of epithelium on fibrocyte survival *in vitro*, macroscopically normal, lung resection material was ethically obtained by lobectomy from a separate group of patients categorized into COPD and control groups as per GOLD criteria (Table E4).

All subjects provided written informed consent to participate to the study. All clinical data were collected in the Clinical Investigation Center (CIC1401) from the University Hospital of Bordeaux. The study protocol was approved by the research ethics committee (“CPP”) and the French National Agency for Medicines and Health Products Safety (“ANSM”).

#### Study design

The study protocol was approved by the local research ethics committee on May 30, 2012 and the French National Agency for Medicines and Health Products Safety on May 22, 2012. All clinical investigations have been conducted according to the principles expressed in the Declaration of Helsinki. All subjects provided written informed consent. The clinical trial was conducted from April 2013 (1^st^ patient, 1^st^ visit) to May 2016 (last patient, last visit). As already indicated, all patients undergoing surgery were thus recruited from the Department of Thoracic Surgery of the University Hospital. The study was sponsored and funded by the University Hospital of Bordeaux (*i.e.* “CHU de Bordeaux”). All authors were academic and made the decision to submit the manuscript for publication and vouch for the accuracy and integrity of the contents. The study has been registered at ClinicalTrials.gov under the N° NCT01692444 (*i.e.* “Fibrochir” study).

The study design is summarized in Fig E1. The pre-inclusion visit (V1) before surgery, consisted of patient information and signature of the inform consent followed by a clinical evaluation (*i.e*., pulmonary auscultation, assessment of the WHO score, history of previous 12 months, smoking status, current treatment…). A full-body CT-scan with injection, was performed as part of the classical disease management but was preceded by two complementary thoracic acquisitions at expiration and inspiration without injection within the framework of the study. The patients also underwent echocardiography and lung function testing using body plethysmography, lung transfer capacity (TLCO) and arterial gas. The inclusion visit (V2), the day of surgery, consisted of a clinical evaluation (*i.e*., control assessment test (CAT), St Georges Quality of Life Questionnaire (SGQLQ) and six-minute walk test) and venous blood sample (50 ml) for fibrocytes analysis. After thoracic surgery (lobectomy or pneumonectomy), a pulmonary sample from a grossly normal part of the surgical specimen is included in paraffin for subsequent analysis of the bronchial fibrocytes. Due to low quality of some tissue sections, fibrocyte density quantification was impossible in 10 distal specimens and 7 proximal specimens (Fig E1), which were excluded from peribronchial fibrocyte analysis. During hospital stay, clinical data was collected such as pTNM status for cancer patients. A visit one month ± 15 days after surgery (V3) consisted of spirometric evaluation. The last visit one year ± 15 days after surgery (V4) consisted of clinical (CAT, SGQLQ and six-minute walk test) and functional (plethysmography, TLCO, arterial gas) evaluations. COPD patients and control subjects performed the whole series of “Fibrochir” visits, with the exception of 2 COPD patients who provided written informed consent for the use of biological samples and clinical data for research and underwent only two visits (corresponding to V1 and V2) including the surgical pulmonary sample for peribronchial fibrocyte analysis.

#### Bronchial fibrocytes identification

A sub-segmental bronchus sample (for proximal tissue) as well as fragments of distal parenchyma were obtained from macroscopically normal lung resection material. The samples were embedded in paraffin and sections of 2.5 µm thick were cut, as described previously (2). Sections were deparaffinized through three changes of xylene and through graded alcohols to water. Heat induced antigen retrieval was performed using citrate buffer, pH 6 (Fisher Scientific, Illkirch, France) in a Pre-Treatment Module (Agilent, Les Ulis, France). Endogenous peroxidase and alkaline phosphatases (AP) were blocked for 10 min using Dual Enzyme Block (Diagomics, Blagnac, France). Nonspecific binding was minimized by incubating the sections with 4% Goat Serum (Agilent) for 30 min. The sections were stained with both rabbit anti-FSP1 polyclonal antibody (Agilent) and mouse anti-CD45 monoclonal antibody (BD Biosciences, San Jose, CA), or mouse anti-CD3 monoclonal antibody (Agilent), mouse anti-CD19 monoclonal (Agilent), mouse anti-CD34 monoclonal antibody (Agilent) or appropriate isotype controls, rabbit IgG (Fisher Scientific) and mouse IgG1 (R&D Systems, Lille, France) at the same concentration. For CD45-FSP1 double staining, the sections were re-incubated with HRP-Polymer anti-Mouse and AP Polymer anti-Rabbit antibodies (Diagomics). Sections were developed with the chromogenic substrates, GBI-Permanent Red and Emerald. For CD3-FSP1, CD19-FSP1 and CD34-FSP1 double staining, the sections were re-incubated with HRP antiMouse (Agilent) and with Alexa488–conjugated anti-Rabbit (Fisher Scientific) antibodies. Immunoreactivity was detected by using the DAB System (Agilent) for CD3, CD19 or CD34 staining and by fluorescence for FSP1 staining.

The sections were imaged using a slide scanner Nanozoomer 2.0HT with fluorescence imaging module (Hamamatsu Photonics, Massy, France) using objective UPS APO 20X NA 0.75 combined to an additional lens 1.75X, leading to a final magnification of 35X. Virtual slides were acquired with a TDI-3CCD camera. Fluorescent acquisitions were done with a mercury lamp (LX2000 200W - Hamamatsu Photonics) and the set of filters adapted for DAPI and Alexa 488. Bright field and fluorescence images where acquired with the NDP-scan software (Hamamatsu) and processed with ImageJ. Quantification of dual positive cells for FSP1 and CD45 was performed, as described in Fig 1. A color deconvolution plugin was used on bright field image to separate channels corresponding to GBI-Permanent Red and Emerald double staining, and a binary threshold was then applied to these images (Fig 1A). Tissue fibrocytes were defined as cells dual positive for cytoplasmic FSP1 and plasma membrane CD45 double staining on the merged threshold image (Fig 1B-C). The lamina propria contour was manually determined on bright field image and the area was calculated. For distal bronchi, the lumen area was also determined and only bronchi less than 2 mm in diameter were analyzed as described by J.C. Hogg *et al* (3). The density of FSP1^+^ CD45^+^ cells was defined by the ratio between the number of dual positive cells in the lamina propria divided by lamina propria area. Quantification of dual positive cells for FSP1 and CD3, FSP1 and CD19 or FSP1 and CD34 was performed, as described above with some modification: a color deconvolution plugin was used on bright field image to select the channel corresponding to DAB signal (for CD3, CD19 or CD34 staining), and a binary threshold was then applied to this image and fluorescence image corresponding to FSP1 staining. Tissue area and cell measurements were all performed in a blinded fashion for patients’ characteristics.

#### Quantitative computed tomography

CT scans were performed on a Somatom Sensation Definition 64 (Siemens, Erlangen, Germany) at full inspiration and expiration, as described previously (4-7). Briefly, quantitative analysis was performed by using dedicated and validated software: Automatic quantification of bronchial wall area (WA), lumen area (LA), WA/LA (WA%) and wall thickness (WT) was obtained on orthogonal bronchial cross sections by using the Laplacian-of-Gaussian algorithm and homemade software (8, 9); Automatic quantification of both emphysema and air trapping was assessed using Myrian software (Intrasense, Montpellier, France) and both low attenuation area per cent (LAA%) (4, 5) and mean lung attenuation (MLA) during expiration (6, 7); Quantification of pulmonary vessels was obtained from CT images, as previously described (4). Briefly, CT set of images reconstructed with sharp algorithm (B70f) were analyzed by using the ImageJ software version 1.40g (a public domain Java image program available at http://rsb.info.nih.gov/ij/). The small pulmonary vessels measurements were made automatically as described elsewhere (4, 10-12). The cross section area (CSA) and cross section number (CSN) of small pulmonary vessels were quantified separately at the subsegmental and at the sub-subsegmental levels (4, 12). The subsegmental and sub-subsegmental levels are defined by a vessel area between 5 and 10 mm^2^ and less than 5 mm^2^, respectively. Finally, quantifications were obtained after normalization by the corresponding lung section area at each CT slice: the cross sectional area of small pulmonary vessel less than 5 mm^2^ (%CSA<5). Four measurements were obtained after normalization by the corresponding lung section area at each CT slice: the cross sectional area of small pulmonary vessel between 5 to 10 mm^2^ (%CSA5-10), and less than 5 mm^2^ (%CSA<5), the mean number of cross-sectioned vessels CSN5-10 and CSN<5.

#### Circulating fibrocytes identification

Non-adherent non-T (NANT) cells were purified from Peripheral Blood Mononuclear Cells (PBMC) separated from the whole blood, and circulating fibrocytes were identified as double positive cells for the surface marker CD45 and the intracellular marker collagen I by flow cytometry, as described previously (13). Briefly, PBMC were first separated from the whole blood by Ficoll-Hypaque (Dutscher, Brumath, France) density gradient centrifugation. The non-adherent mononuclear cell fraction was taken and washed in cold PBS containing 0.5% bovine serum albumin (BSA, Sigma-Aldrich) and 2 mM Ethylene Diamine Tetra-acetic Acid (EDTA, Invitrogen). T-cells were further depleted with anti-CD3 monoclonal antibody (Miltenyi Biotech, Paris, France). Cells were fixed overnight with Cytofix/Cytoperm (eBioscience, Paris, France), washed in permeabilization buffer (eBioscience) and incubated either with mouse anti-human collagen I antibody (Millipore, St-Quentin-en-Yvelines, France) or with matched IgG1 isotype control (Santa Cruz Biotechnology, Heidelberg, Germany), followed by fluorescein isothiocyanate (FITC)–conjugated anti-mouse antibodies (Beckman Coulter, Villepinte, France). Next, the cell pellet was incubated either with allophycocyanin (APC)-conjugated anti-CD45 antibodies (BD Biosciences, San Jose, CA) or with matched APC-conjugated IgG1 isotype control (BD Biosciences). The cell suspension was analyzed with a BD FACSCanto II flow cytometer (BD Biosciences). Offline analysis was performed with FACSDiva (BD Biosciences) and FlowJo (Tree Star, Ashland, OR) software. The negative threshold for CD45 was set using a matched APC-conjugated IgG1 isotype control, and all subsequent samples were gated for the CD45 positive region. Cells gated for CD45 were analyzed for collagen I expression, with negative control thresholds set using FITC-stained cells. Specific staining for collagen I was determined as an increase in positive events over this threshold. Fibrocytes numbers were expressed as both a percentage of total PBMC counts and as absolute number of cells.

#### Bronchial epithelial supernatants

Human bronchial epithelial cells (BEC) were derived from bronchial specimens as described previously (14). Bronchial epithelial tissue was cultured in bronchial epithelial growth medium (Stemcell, Grenoble, France) in a flask (0.75 cm^2^). After confluence, basal BEC were plated (2.10^5^ cells per well) on uncoated nucleopore membranes (24-mm diameter, 0.4-µm pore size, Transwell Clear; Costar, Cambridge, Mass) in ALI medium (Stemcell) applied at the basal side only to establish the air-liquid interface. Cells were maintained in culture for 21 days to obtain a differentiated cell population with a mucociliary phenotype. Basal epithelial supernatant was collected every 2-3 days and used for further experiments.

#### Fibrocyte differentiation and survival

A total of 2.10^6^ NANT cells resuspended in 0.2 ml DMEM (Fisher Scientific), containing 4.5 g/l glucose and glutaMAX, supplemented with 20% fetal calf serum (Biowest, Riverside, USA), penicillin/streptomycin and MEM non-essential amino acid solution (Sigma-Aldrich), was added to each well of a 6 well plate. After one week in culture, fibrocyte differentiation was induced by changing the medium for a serum-free medium (Fig E2), or for a serum-free medium containing 50% of basal epithelium supernatant (Fig E10). Mediums were changed every 2-3 days. After 2 weeks in culture, the cells were detached by accutase treatment (Fisher Scientific), fixed overnight with Cytofix/Cytoperm and washed in permeabilization buffer. Cells were incubated either with rabbit anti-FSP1 polyclonal antibody (Agilent) or with matched IgG isotype control (Fisher Scientific), followed by Phycoerythrin (PE)–conjugated anti-rabbit antibody (Santa Cruz Biotechnology, Heidelberg, Germany). Next, the cell pellet was incubated either with FITC–conjugated mouse anti-human collagen I antibody (Millipore) or with matched FITC-conjugated IgG1 isotype control (Millipore) and with APC-conjugated anti-CD45 antibodies or with matched APC-conjugated IgG1 isotype control. The cell suspension was analyzed with a BD FACSCanto II flow cytometer. Cells gated for CD45 and FSP1 were analyzed for collagen-1 expression, with negative control thresholds set using isotype-stained cells. Specific staining for collagen-1 was determined as an increase in positive events over this threshold.

Propidium iodide (Fisher Scientific) was used for the detection of dying cells. After 2 weeks in culture, cells detached by accutase treatment were used to prepare single cell suspension at 1.10^6^ cells/ml. After addition of propidium iodide, the dying cells (PI^+^ cells) were detected by flow cytometry (BD Biosciences).

#### Statistical analysis

Primary outcome was the density of bronchial fibrocytes in both distal and proximal airways. Secondary outcomes were lung function parameters, CT parameters and the percentage of blood fibrocytes in PBMC. The statistical analysis was performed with Prism 6 software (GraphPad, La Jolla, CA) and NCSS software (NCSS 2001, Kaysville, UT, USA). Values are presented as medians with individual plots or means ± SD. Statistical significance, defined as *P* < 0.05, was analyzed by Fisher’s exact tests for comparison of proportions, by two-sided independent t-tests for variables with a parametric distribution, and, by Mann–Whitney U tests, Wilcoxon tests and Spearman correlation coefficients for variables with a non-parametric distribution. Receiver operating characteristic (ROC) curves were built with NCSS software (NCSS 2001, Kaysville, UT, USA) and ROC analysis was performed to determine areas under the curve (AUC) and cut-off values for the best fibrocytes density in distal and proximal tissue specimens to predict COPD. Those 2 cut-off values were then used to evaluate the association between COPD and a high density of tissue fibrocytes using a univariate logistic regression analysis.

## Supplemental tables

**Table E1.**
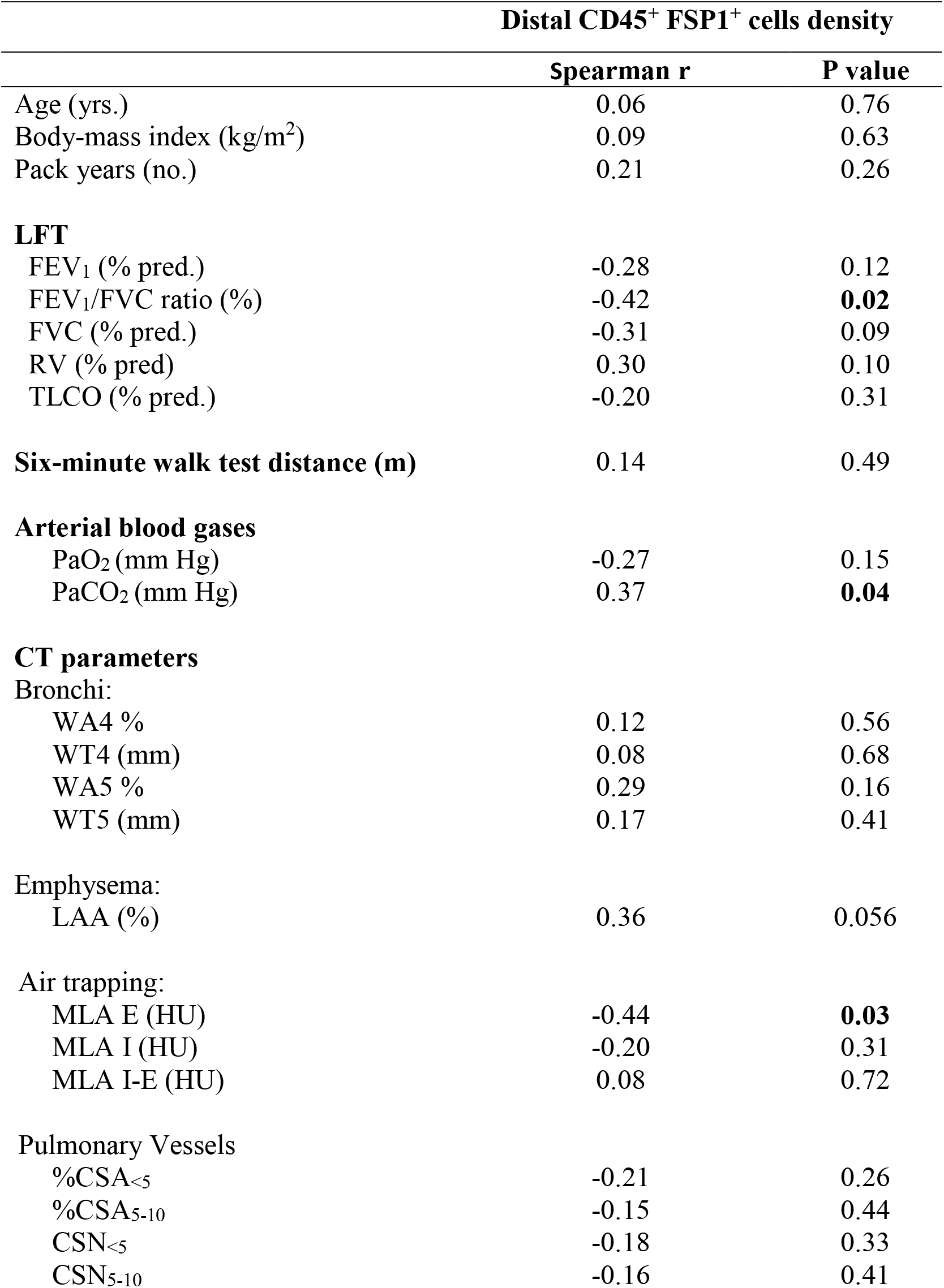
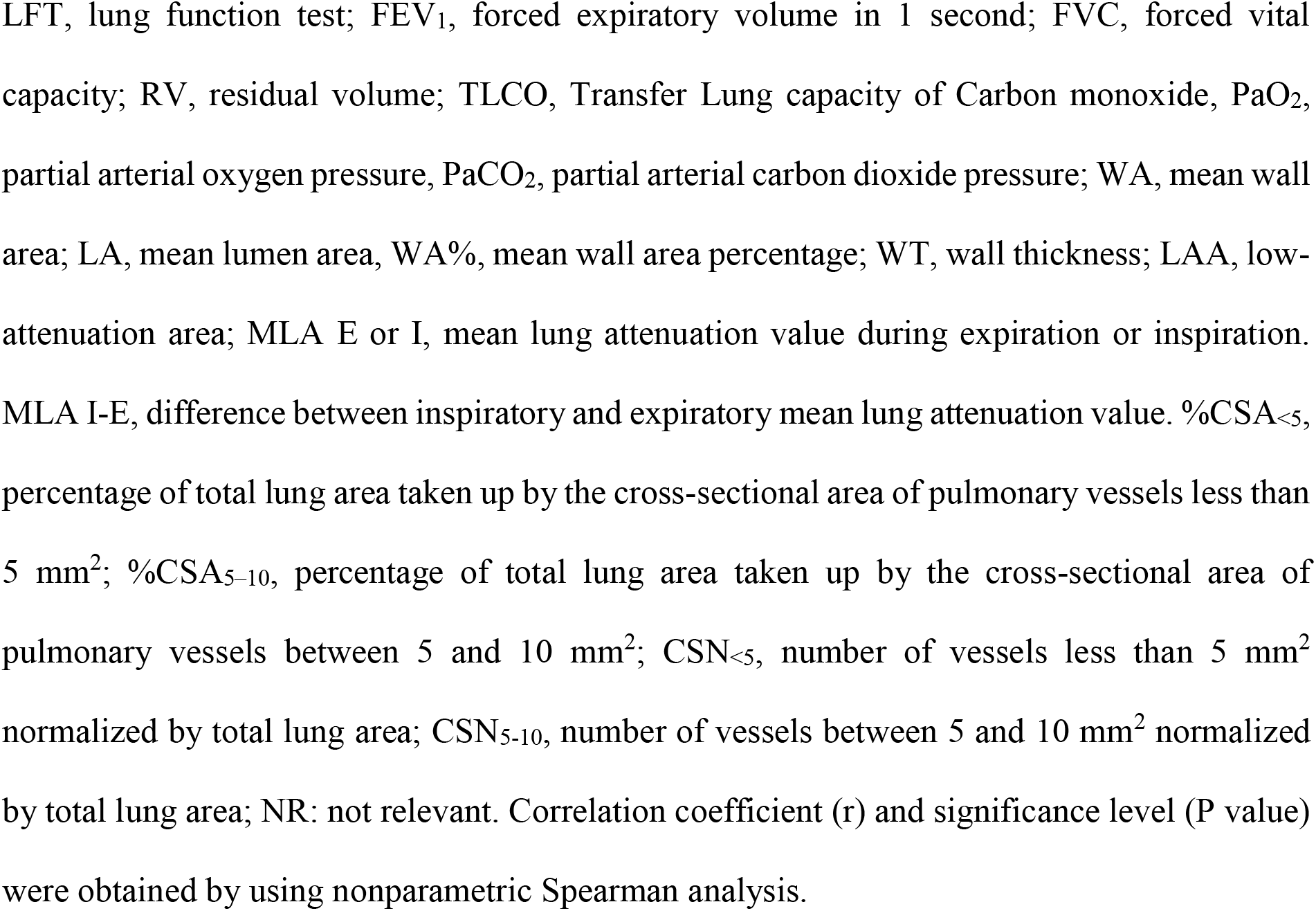
Association between distributions of distal tissue fibrocytes and COPD clinical characteristics

|  | Distal CD45<sup>+</sup> FSP1<sup>+</sup> cells density |  |
| --- | --- | --- |
|  | Spearman r | P value |
| Age (yrs.) | 0.06 | 0.76 |
| Body-mass index (kg/m<sup>2</sup>) | 0.09 | 0.63 |
| Pack years (no.) | 0.21 | 0.26 |
| **LFT** |  |  |
| FEV<sub>1</sub> (% pred.) | -0.28 | 0.12 |
| FEV<sub>1</sub>/FVC ratio (%) | -0.42 | **0.02** |
| FVC (% pred.) | -0.31 | 0.09 |
| RV (% pred) | 0.30 | 0.10 |
| TLCO (% pred.) | -0.20 | 0.31 |
| **Six-minute walk test distance (m)** | 0.14 | 0.49 |
| **Arterial blood gases** |  |  |
| PaO<sub>2</sub> (mm Hg) | -0.27 | 0.15 |
| PaCO<sub>2</sub> (mm Hg) | 0.37 | **0.04** |
| **CT parameters** |  |  |
| Bronchi: |  |  |
| WA4 % | 0.12 | 0.56 |
| WT4 (mm) | 0.08 | 0.68 |
| WA5 % | 0.29 | 0.16 |
| WT5 (mm) | 0.17 | 0.41 |
| Emphysema: |  |  |
| LAA (%) | 0.36 | 0.056 |
| Air trapping: |  |  |
| MLA E (HU) | -0.44 | **0.03** |
| MLA I (HU) | -0.20 | 0.31 |
| MLA I-E (HU) | 0.08 | 0.72 |
| Pulmonary Vessels |  |  |
| %CSA<sub><5</sub> | -0.21 | 0.26 |
| %CSA<sub>5-10</sub> | -0.15 | 0.44 |
| CSN<sub><5</sub> | -0.18 | 0.33 |
| CSN<sub>5-10</sub> | -0.16 | 0.41 |

LFT, lung function test; FEV<sub>1</sub>, forced expiratory volume in 1 second; FVC, forced vital capacity; RV, residual volume; TLCO, Transfer Lung capacity of Carbon monoxide, PaO<sub>2</sub>, partial arterial oxygen pressure, PaCO<sub>2</sub>, partial arterial carbon dioxide pressure; WA, mean wall area; LA, mean lumen area, WA%, mean wall area percentage; WT, wall thickness; LAA, low-attenuation area; MLA E or I, mean lung attenuation value during expiration or inspiration. MLA I-E, difference between inspiratory and expiratory mean lung attenuation value. %CSA<sub><5</sub>, percentage of total lung area taken up by the cross-sectional area of pulmonary vessels less than 5 mm<sup>2</sup>; %CSA<sub>5-10</sub>, percentage of total lung area taken up by the cross-sectional area of pulmonary vessels between 5 and 10 mm<sup>2</sup>; CSN<sub><5</sub>, number of vessels less than 5 mm<sup>2</sup> normalized by total lung area; CSN<sub>5-10</sub>, number of vessels between 5 and 10 mm<sup>2</sup> normalized by total lung area; NR: not relevant. Correlation coefficient (r) and significance level (P value) were obtained by using nonparametric Spearman analysis.

**Table E2.**
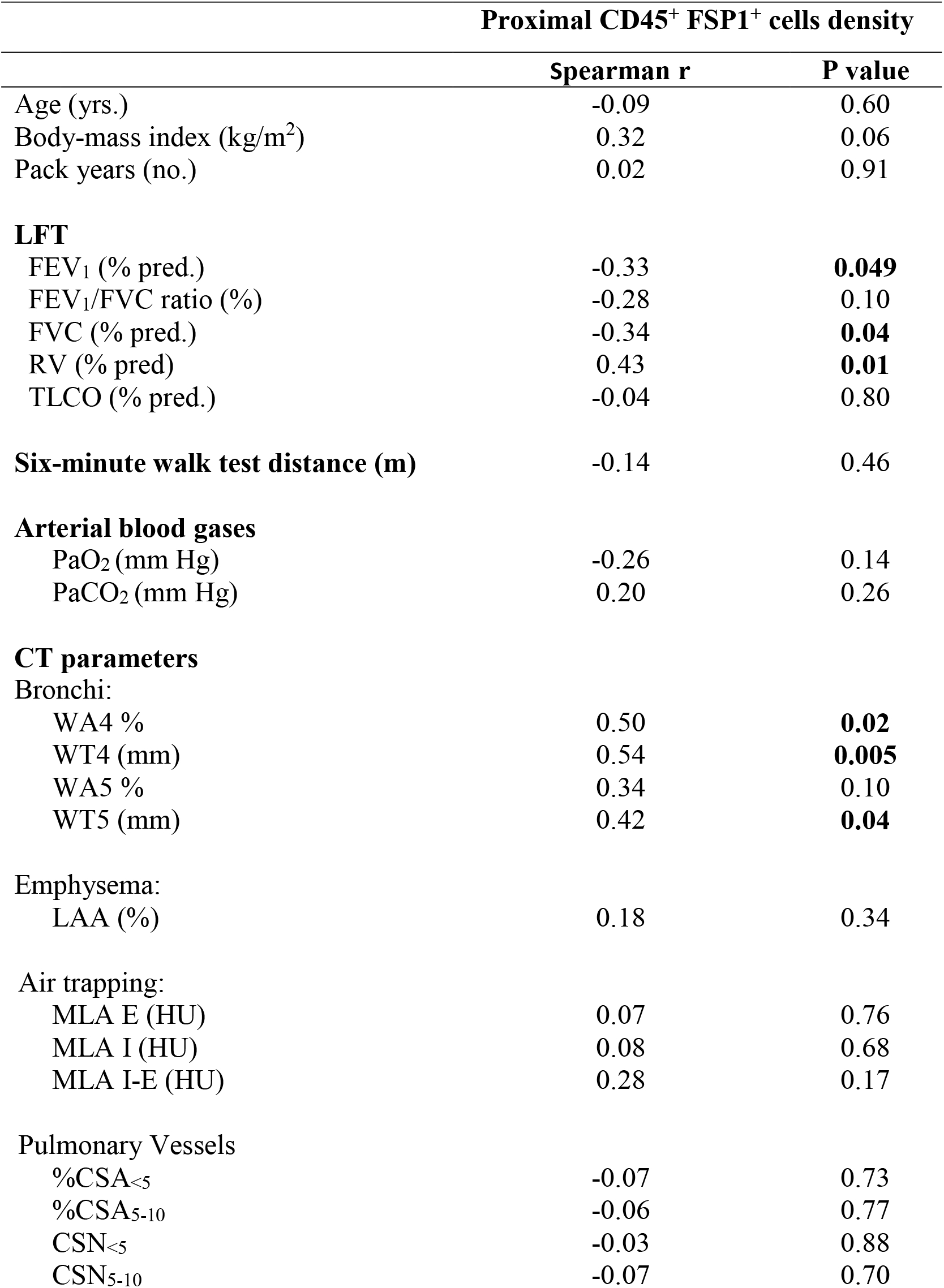
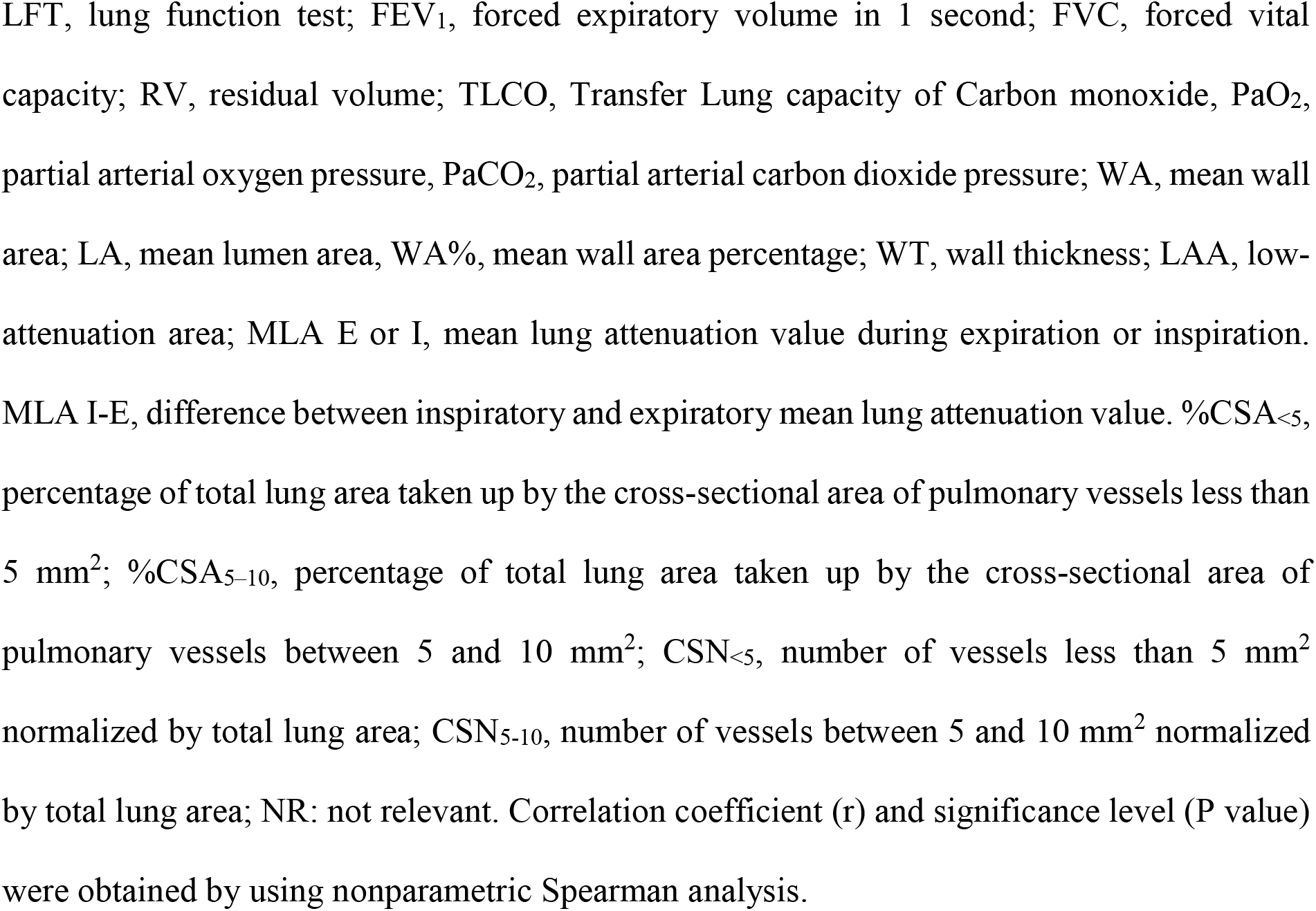
Association between distributions of proximal tissue fibrocytes and COPD clinical characteristics

|  | Proximal CD45<sup>+</sup> FSP1<sup>+</sup> cells density |  |
| --- | --- | --- |
|  | Spearman r | P value |
| Age (yrs.) | -0.09 | 0.60 |
| Body-mass index (kg/m<sup>2</sup>) | 0.32 | 0.06 |
| Pack years (no.) | 0.02 | 0.91 |
| **LFT** |  |  |
| FEV<sub>1</sub> (% pred.) | -0.33 | **0.049** |
| FEV<sub>1</sub>/FVC ratio (%) | -0.28 | 0.10 |
| FVC (% pred.) | -0.34 | **0.04** |
| RV (% pred) | 0.43 | **0.01** |
| TLCO (% pred.) | -0.04 | 0.80 |
| **Six-minute walk test distance (m)** | -0.14 | 0.46 |
| **Arterial blood gases** |  |  |
| PaO<sub>2</sub> (mm Hg) | -0.26 | 0.14 |
| PaCO<sub>2</sub> (mm Hg) | 0.20 | 0.26 |
| **CT parameters** |  |  |
| Bronchi: |  |  |
| WA4 % | 0.50 | **0.02** |
| WT4 (mm) | 0.54 | **0.005** |
| WA5 % | 0.34 | 0.10 |
| WT5 (mm) | 0.42 | **0.04** |
| Emphysema: |  |  |
| LAA (%) | 0.18 | 0.34 |
| Air trapping: |  |  |
| MLA E (HU) | 0.07 | 0.76 |
| MLA I (HU) | 0.08 | 0.68 |
| MLA I-E (HU) | 0.28 | 0.17 |
| Pulmonary Vessels |  |  |
| %CSA<sub><5</sub> | -0.07 | 0.73 |
| %CSA<sub>5-10</sub> | -0.06 | 0.77 |
| CSN<sub><5</sub> | -0.03 | 0.88 |
| CSN<sub>5-10</sub> | -0.07 | 0.70 |

LFT, lung function test; FEV<sub>1</sub>, forced expiratory volume in 1 second; FVC, forced vital capacity; RV, residual volume; TLCO, Transfer Lung capacity of Carbon monoxide, PaO<sub>2</sub>, partial arterial oxygen pressure, PaCO<sub>2</sub>, partial arterial carbon dioxide pressure; WA, mean wall area; LA, mean lumen area, WA%, mean wall area percentage; WT, wall thickness; LAA, low-attenuation area; MLA E or I, mean lung attenuation value during expiration or inspiration. MLA I-E, difference between inspiratory and expiratory mean lung attenuation value. %CSA<sub><5</sub>, percentage of total lung area taken up by the cross-sectional area of pulmonary vessels less than 5 mm<sup>2</sup>; %CSA<sub>5-10</sub>, percentage of total lung area taken up by the cross-sectional area of pulmonary vessels between 5 and 10 mm<sup>2</sup>; CSN<sub><5</sub>, number of vessels less than 5 mm<sup>2</sup> normalized by total lung area; CSN<sub>5-10</sub>, number of vessels between 5 and 10 mm<sup>2</sup> normalized by total lung area; NR: not relevant. Correlation coefficient (r) and significance level (P value) were obtained by using nonparametric Spearman analysis.

**Table E3.** Patient characteristics

|  | **COPD** |
| --- | --- |
| n | 3 |
| Age (yr) | 68.7 ± 15.0 |
| Sex (Men/Woman) | 2/1 |
| Body-mass index (kg/m<sup>2</sup>) | 21.0 ± 3.5 |
| Current smoker (Y/N) | 2/1 |
| Former smoker (Y/N) | 1/2 |
| Pack years (no.) | 42.3 ± 24.2 |
| **PFT** |  |
| FEV<sub>1</sub> (% pred.) | 58.3 ± 22.5 |
| FEV<sub>1</sub>/FVC ratio (%) | 53.3 ± 11.5 |
| FVC (% pred.) | 84.0 ± 27.6 |
| **Six-minute walk test distance (m)** | 550 ± 42 |
| **Arterial blood gases** |  |
| PaO<sub>2</sub> (mm Hg) | 77.0 ± 6.1 |
| PaCO<sub>2</sub> (mm Hg) | 37.0 ± 4.2 |

Plus-minus values are means ± SD. PFT, pulmonary function test; FEV<sub>1</sub>, forced expiratory volume in 1 second; FVC, forced vital capacity; PaO<sub>2</sub>, partial arterial oxygen pressure, PaCO<sub>2</sub>, partial arterial carbon dioxide pressure.

**Table E4.** Patient characteristics (for bronchial epithelial supernatants production)

|  | **COPD** | **Control** |
| --- | --- | --- |
| n | 2 | 2 |
| Age (yr) | 62.5 ± 12.0 | 70.5 ± 13.4 |
| Sex (Men/Woman) | 0/2 | 2/0 |
| Body-mass index (kg/m<sup>2</sup>) | 31.5 ± 7.8 | 24.5 ± 0.7 |
| Current smoker (Y/N) | 0/1* | 0/2 |
| Former smoker (Y/N) | 1/0* | 0/2 |
| Pack years (no.) | 50.0 ± 70.7 | 0 |
| **PFT** |  |  |
| FEV<sub>1</sub> (% pred.) | 62.5 ± 3.5 | 113.5 ± 62.9 |
| FEV<sub>1</sub>/FVC ratio (%) | 61.0 ± 0 | 81.5 ± 0.1 |

Plus-minus values are means ± SD. PFT, pulmonary function test; FEV<sub>1</sub>, forced expiratory volume in 1 second; FVC, forced vital capacity. \* one of the two COPD patients has been professionally exposed to noxious particles.

**Table E5.** Patient characteristics

|  | **COPD** |
| --- | --- |
| n | 6 |
| Age (yr) | 69.2 ± 8.0 |
| Sex (Men/Woman) | 2/4 |
| Body-mass index (kg/m<sup>2</sup>) | 27.7 ± 5.6 |
| Current smoker (Y/N) | 4/2 |
| Former smoker (Y/N) | 2/4 |
| Pack years (no.) | 66.3 ± 24.7 |
| **PFT** |  |
| FEV<sub>1</sub> (% pred.) | 67.1 ± 22.5 |
| FEV<sub>1</sub>/FVC ratio (%) | 62.1 ± 6.4 |
| FVC (% pred.) | 86.2 ± 30.0 |
| **Six-minute walk test distance (m)** | 395 ± 187 |
| **Arterial blood gases** |  |
| PaO<sub>2</sub> (mm Hg) | 71.1 ± 6.6 |
| PaCO<sub>2</sub> (mm Hg) | 37.1 ± 4.7 |

Plus-minus values are means ± SD. PFT, pulmonary function test; FEV<sub>1</sub>, forced expiratory volume in 1 second; FVC, forced vital capacity; PaO<sub>2</sub>, partial arterial oxygen pressure, PaCO<sub>2</sub>, partial arterial carbon dioxide pressure.

**Fig E1.**
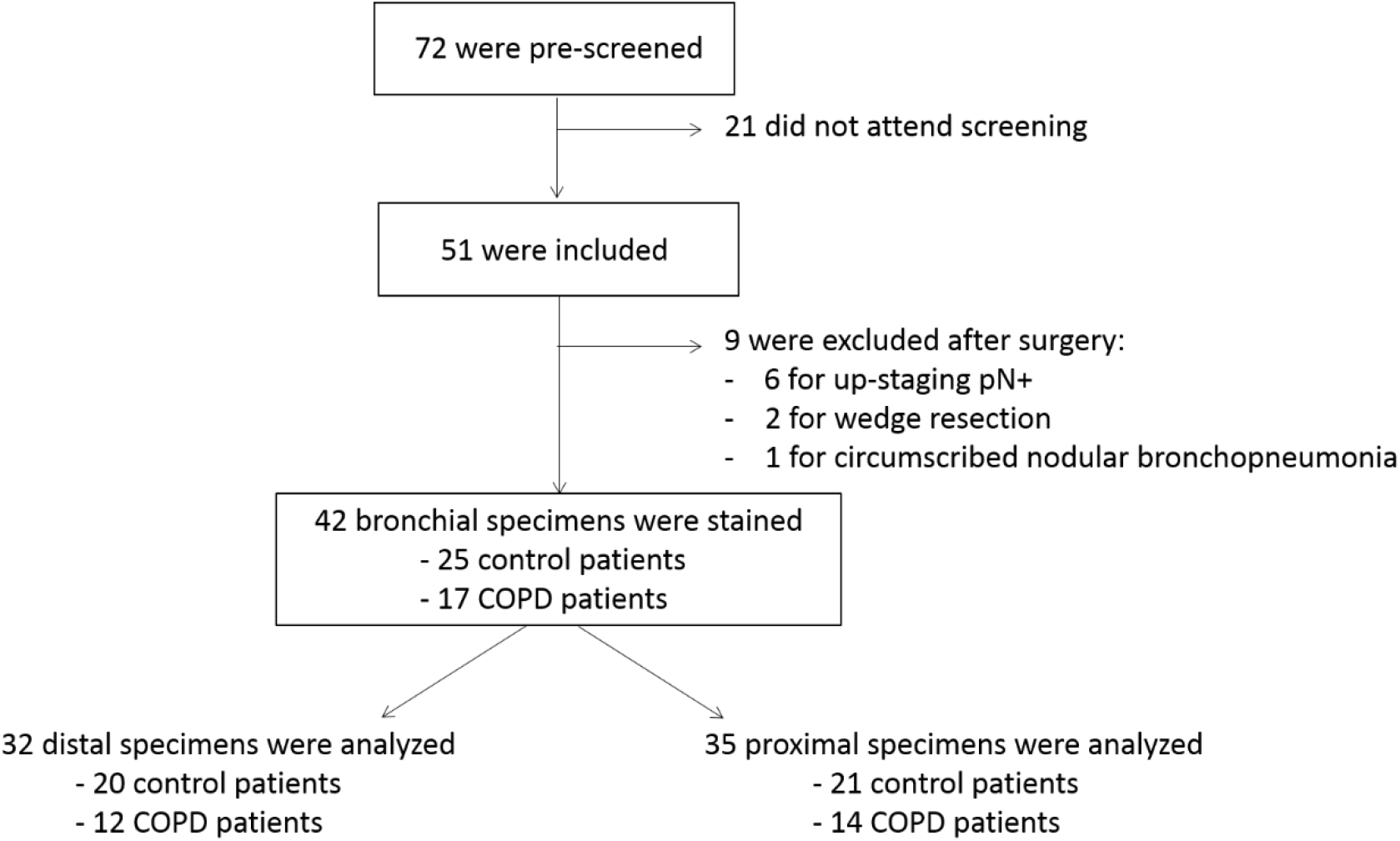
Study design. Numbers of patients who were included and had fibrocyte density quantification in proximal and distal airways.

**Fig E2.**
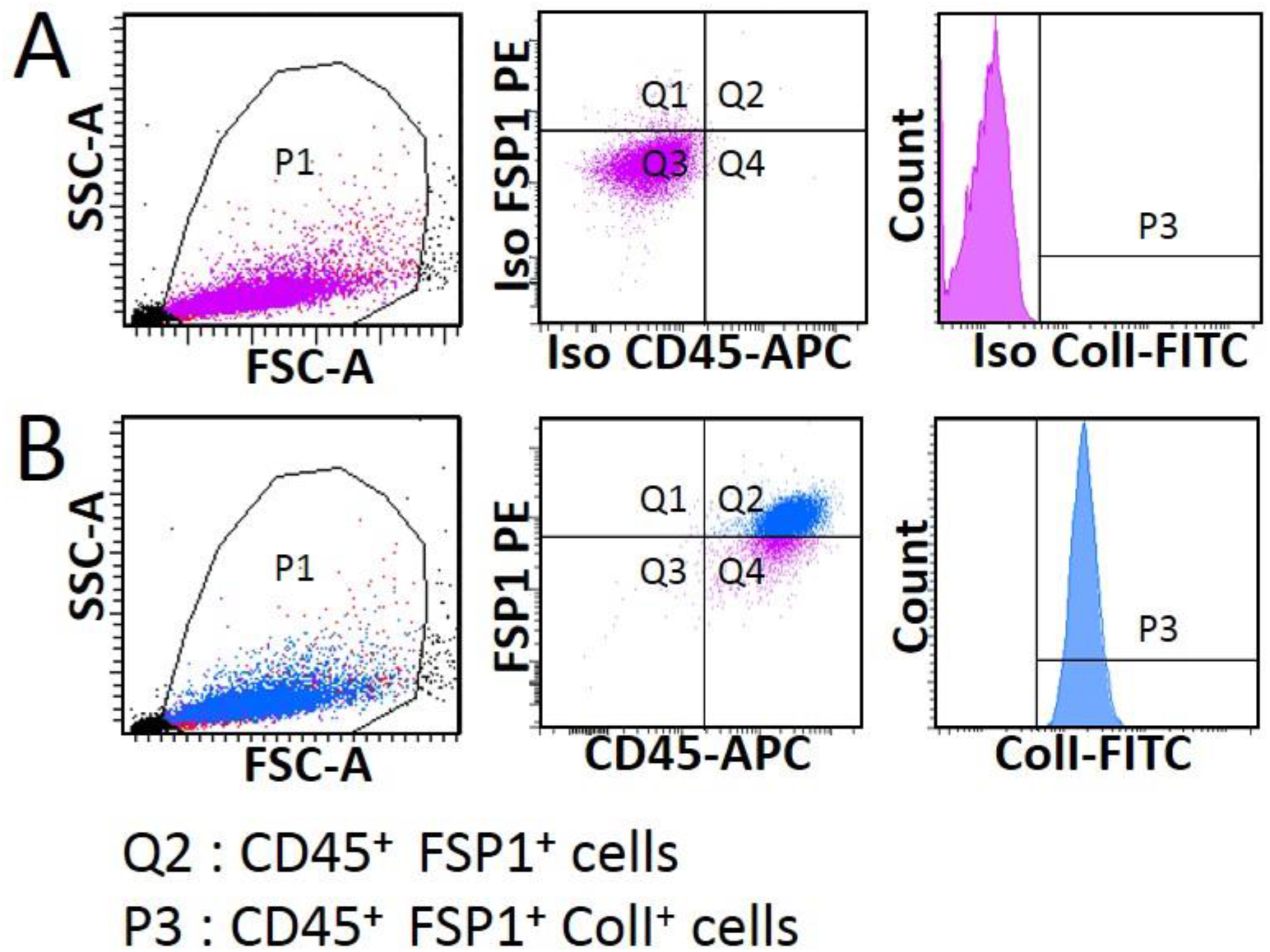
CD45^+^ FSP1^+^ cells purified from NANT cells express collagen I after 2 weeks of differentiation in culture. Representative dot plots of flow cytometry for CD45, FSP1 and collagen I expression. Left panels: total Non-Adherent Non T (NANT) cell population selected on the scatter plot of FSC-A vs SSC-A. Middle panels: a gate (CD45^+^ and FSP1^+^) was drawn in FSP1-PE vs APC-CD45 dot plot to define the Q2 population (positive population for both CD45 and FSP1). Right panels: a gate (collagen I^+^) was drawn in FITC-collagen I (colI) histogram to define the P3 population (positive population for CD45, FSP1 and collagen I). A, isotype control for CD45, FSP1 and collagen I. B, CD45, FSP1 and collagen I stainings. APC: allophycocyanin; FITC: fluorescein isothiocyanate; PE: Phycoerythrin; FSC-A: forward scatter; SSC-A: side scatter.

**Fig E3.**
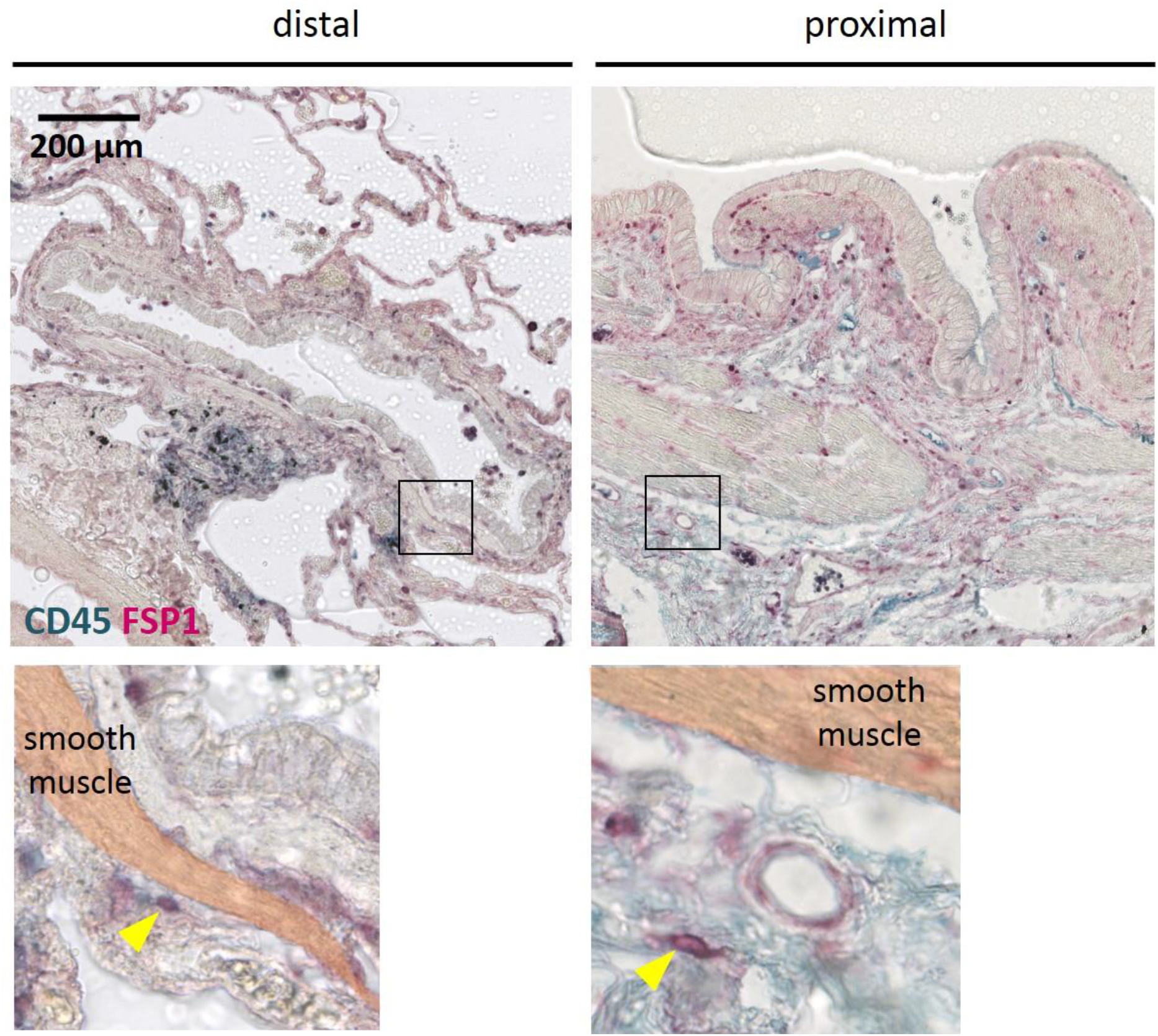
Presence of fibrocytes in peribronchial area outside the smooth muscle layer. Representative staining of CD45 (green) and FSP1 (red) in distal (left) and proximal (right) bronchial tissue specimens. The lower panels show higher magnification of the small area (black boxes) defined in the upper panels. The smooth muscle layer has been highlighted in orange. The yellow arrowheads indicate fibrocytes, defined as CD45^+^ FSP1^+^ cells.

**Fig E4.**
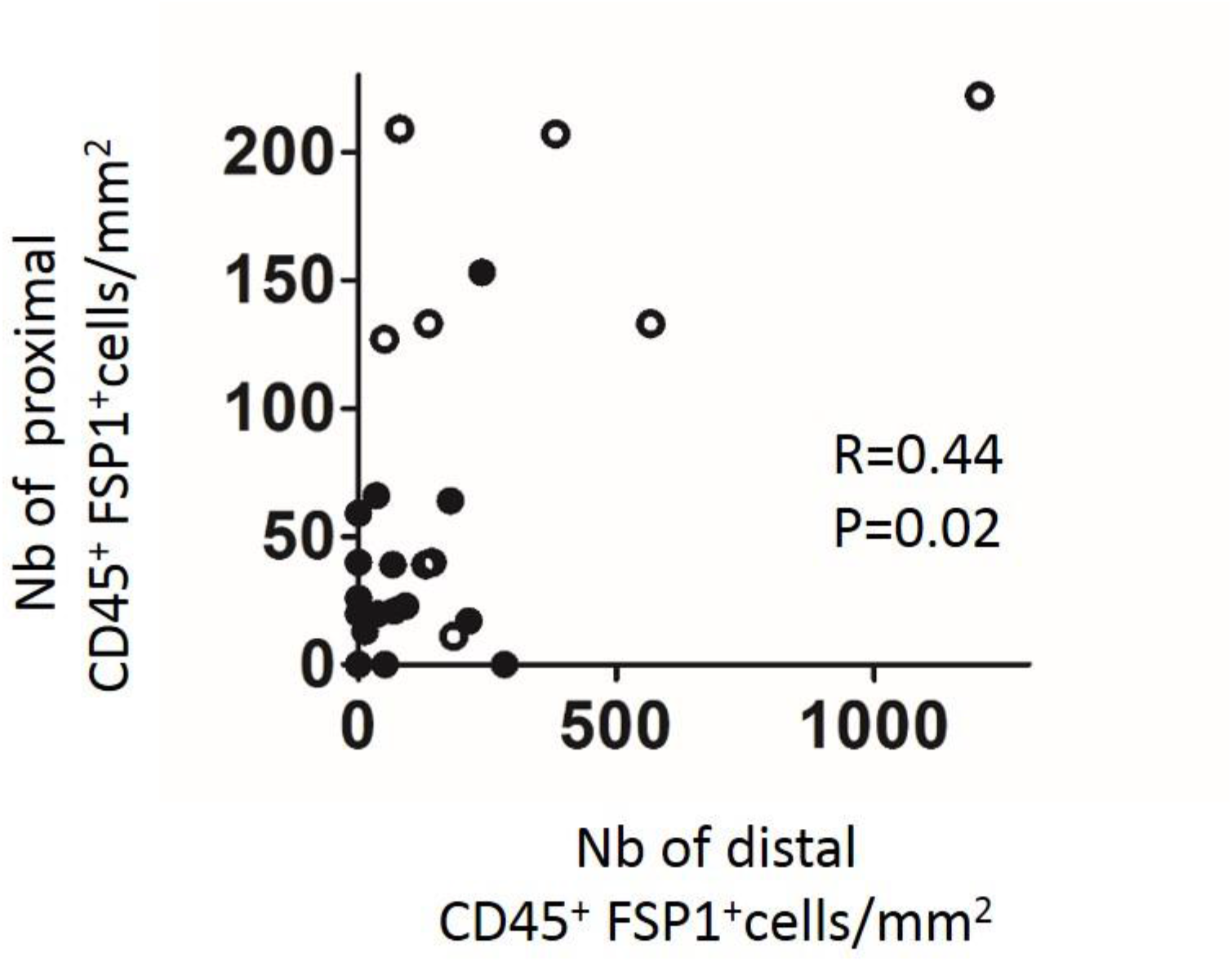
Supplementary Fig E4. Relationship between the density of CD45^+^ FSP1^+^ cells in proximal airways and that in distal airways. Densities measured in control subjects and COPD patients are represented respectively with black and open circles. Correlation coefficient (r) and significance level (P value) were obtained by using nonparametric Spearman analysis.

**Fig E5.**
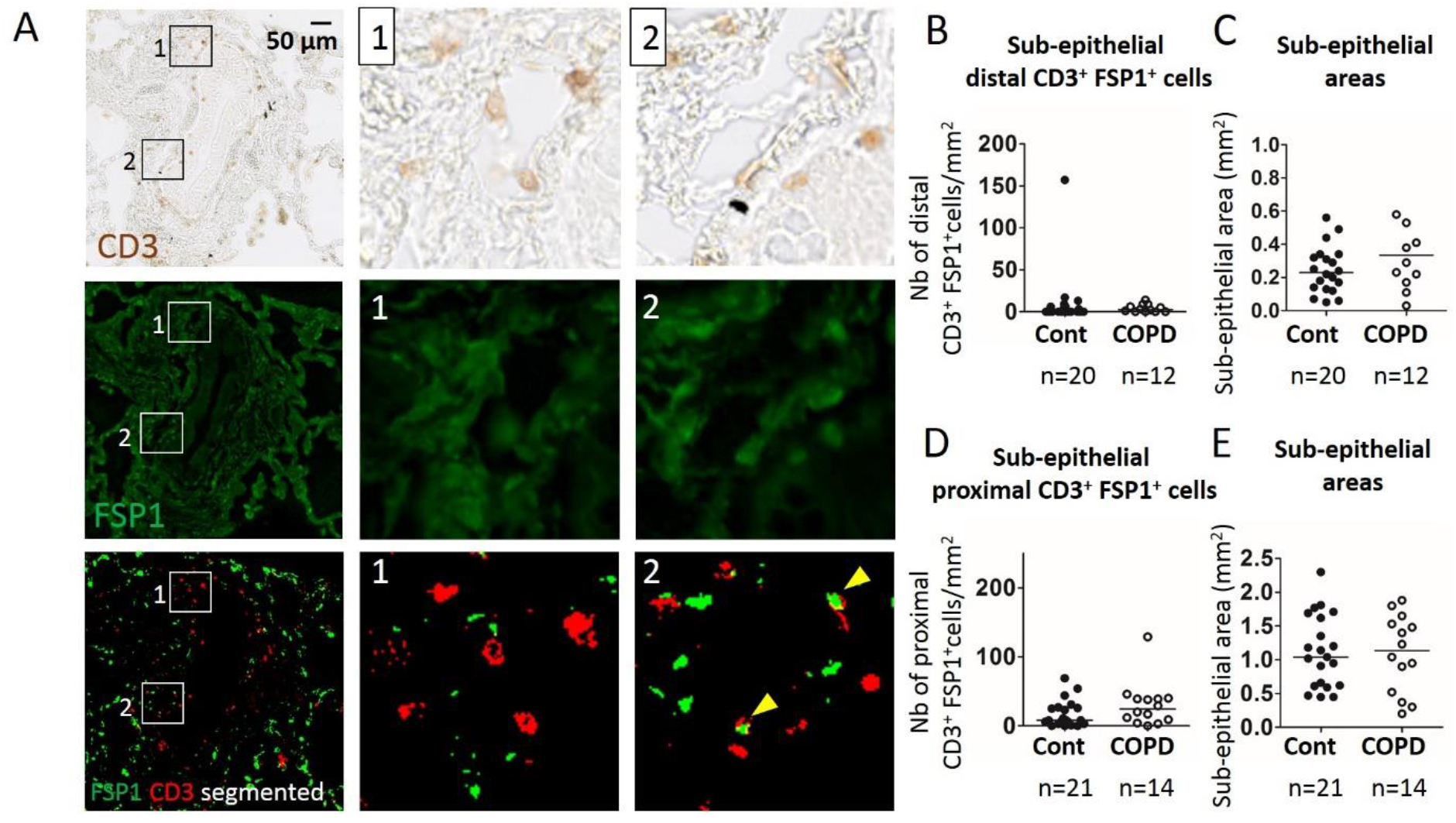
CD3^+^ FSP1^+^ cells represent a minor fraction of CD45^+^ FSP1^+^ cells. A, Representative stainings of CD3 (brown, top panels) and FSP1 (green, middle panels) and merged segmented images (CD3, red and FSP1, green, bottom panels) in distal lung tissue from COPD patient. Middle and right columns represent higher magnification of images in the left column. The yellow arrowheads indicate CD3^+^ FSP1^+^ cells. B, D, Quantification of CD3^+^ FSP1^+^ cells density (normalized by the sub-epithelial area) in distal (B) and proximal (D) tissue specimens from control subjects and COPD patients. C, E, Comparison of sub-epithelial areas in distal (C) and proximal (E) tissue specimens from control subjects and COPD patients. B-D, medians are represented as horizontal lines.

**Fig E6.**
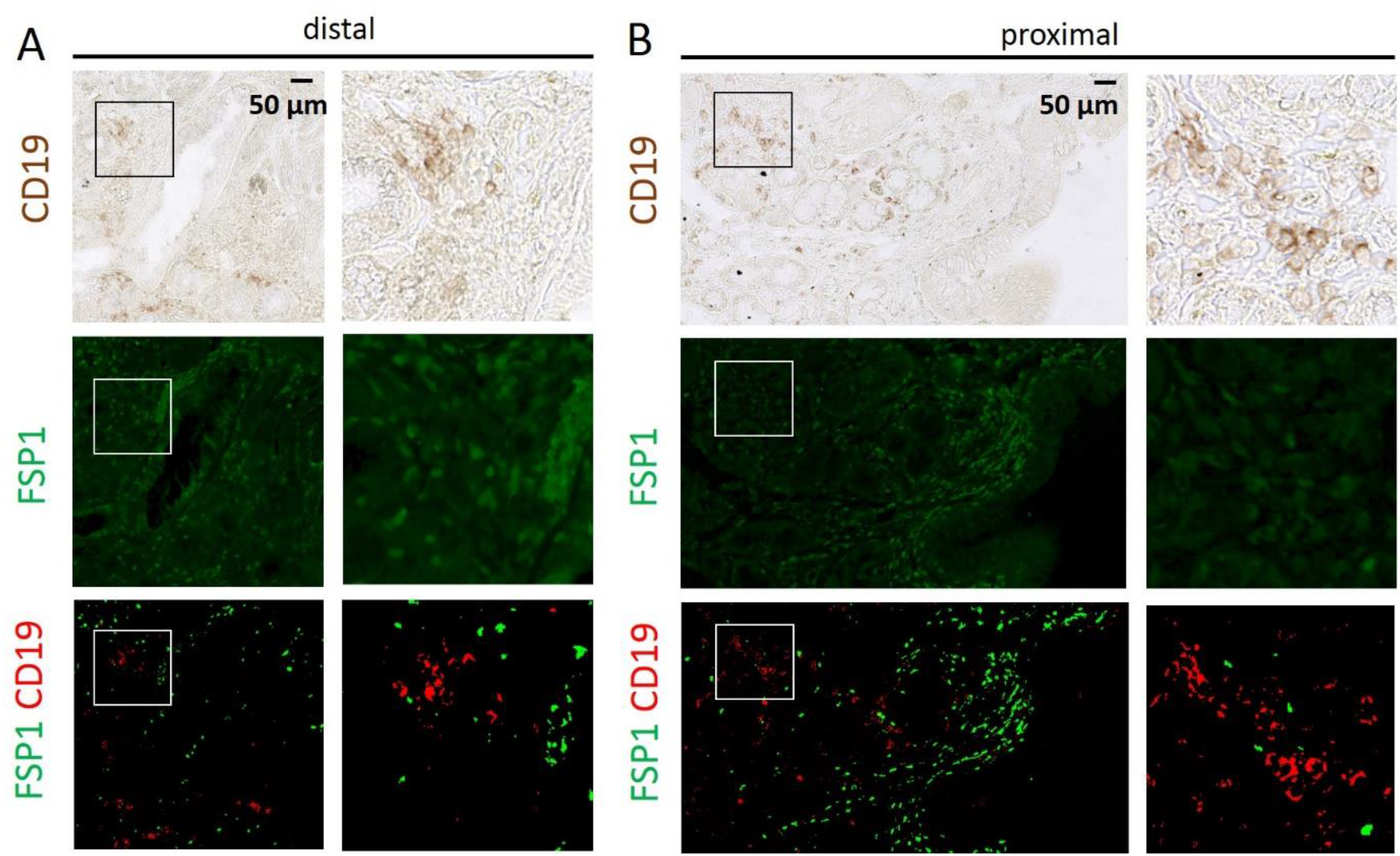
CD19^+^ FSP1^+^ cells are detected neither in proximal and nor in distal airways. A-B, Representative stainings of CD19 (brown, top panels) and FSP1 (green, middle panels) and merged segmented images (CD19, red and FSP1, green, bottom panels) in distal (A) and proximal (B) bronchial tissue specimens from COPD patient. The right columns represent higher magnification of images in the left columns.

**Fig E7.**
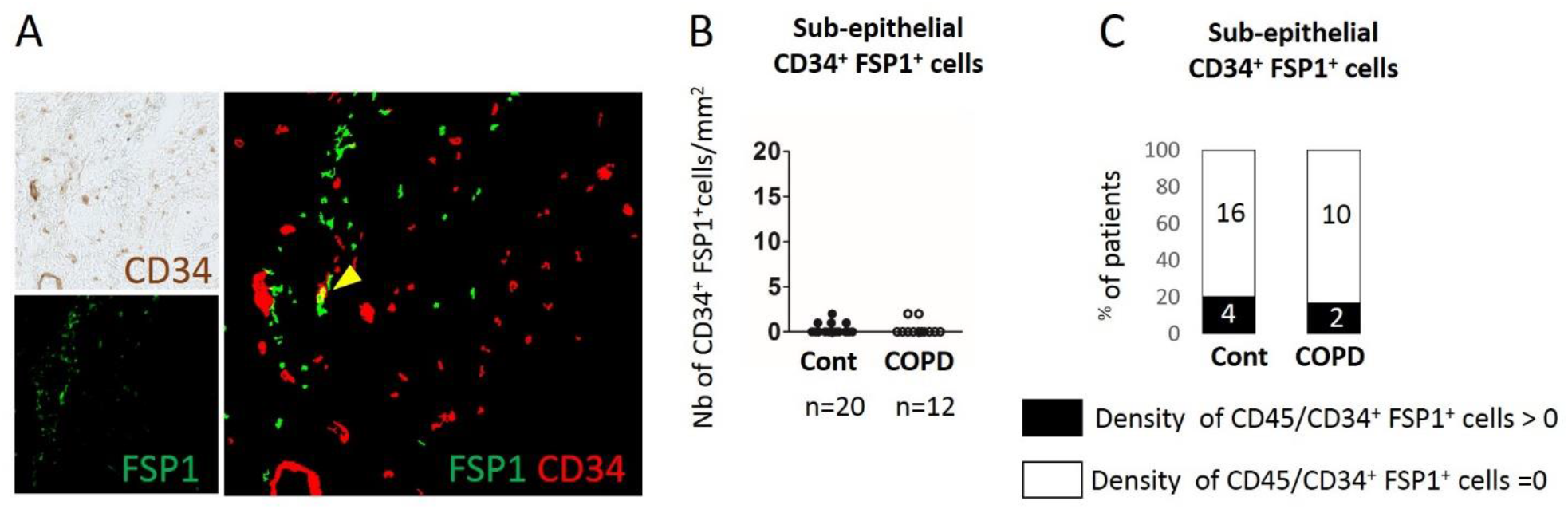
CD34^+^ FSP1^+^ cells are almost absent in distal airways. A, Representative stainings of CD34 (brown, top left panel) and FSP1 (green, bottom left panels) and merged segmented image (CD34, red and FSP1, green, right panel) in distal tissue specimen from COPD patient. The yellow arrowhead indicates CD34^+^ FSP1^+^ cells. B, Quantification of CD34^+^ FSP1^+^ cells density (normalized by the sub-epithelial area) in distal tissue specimens from control subjects and COPD patients. Median is represented as horizontal line. C, Percentage of tissue specimens in which CD34^+^ FSP1^+^ cells have been detected (density>0, black bars) or undetected (density=0, white bars) in distal tissue specimens from control subjects and COPD patients.

**Fig E8.**
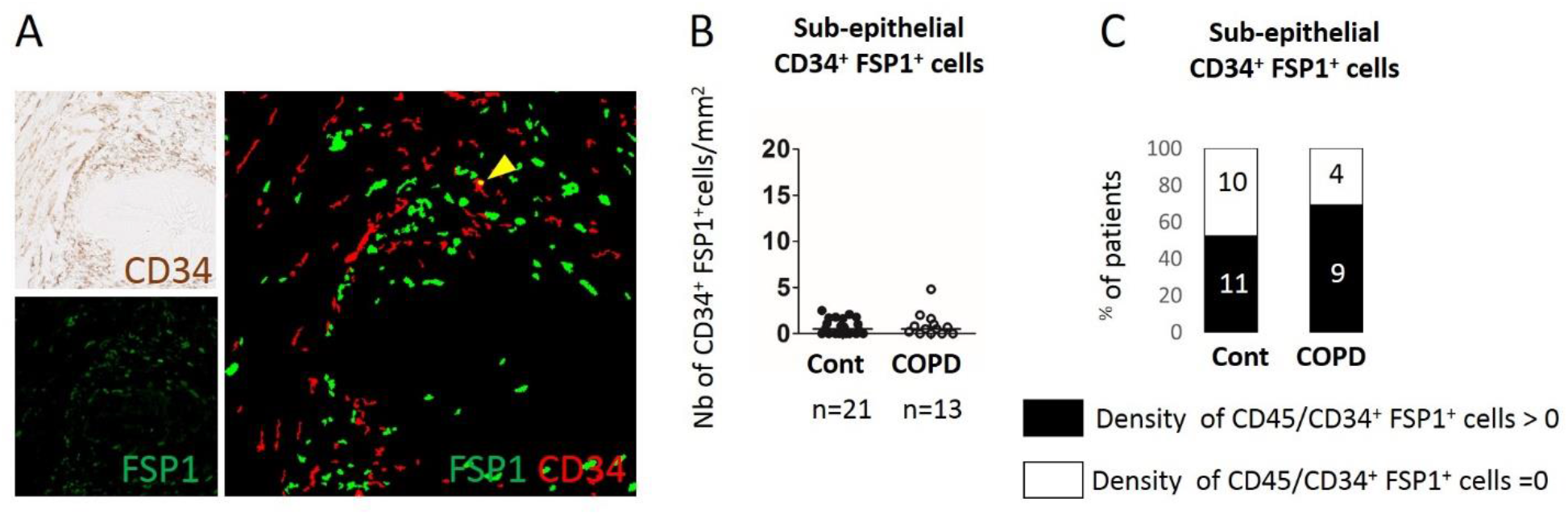
CD34^+^ FSP1^+^ cells are present at a very low level in proximal airways. A, Representative stainings of CD34 (brown, top left panel) and FSP1 (green, bottom left panels) and merged segmented image (CD34, red and FSP1, green, right panel) in proximal tissue specimen from COPD patient. The yellow arrowhead indicates CD34^+^ FSP1^+^ cells. B, Quantification of CD34^+^ FSP1^+^ cells density (normalized by the sub-epithelial area) in proximal tissue specimens from control subjects and COPD patients. Median is represented as horizontal line. C, Percentage of tissue specimens in which CD34^+^ FSP1^+^ cells have been detected (density>0, black bars) or undetected (density=0, white bars) in proximal tissue specimens from control subjects and COPD patients.

**Fig E9.**
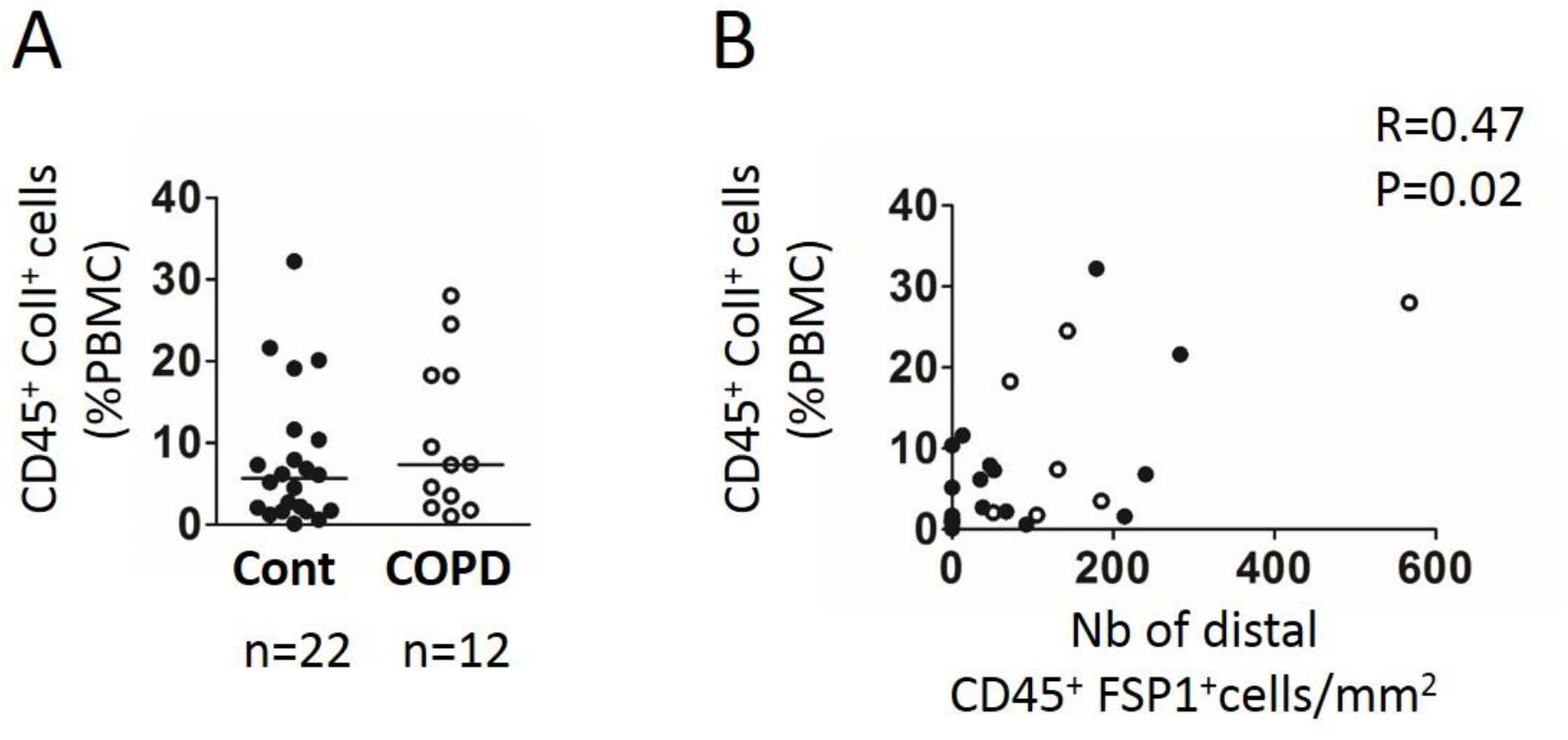
Level of circulating fibrocytes and relationship with tissue fibrocytes density. A, Level of circulating fibrocytes (CD45^+^ ColI^+^ cells), expressed as percentage of PBMC, measured in blood from control subjects (“Cont”, n = 22), and COPD patients (“COPD”, n=12). Medians are represented as horizontal lines. B, Relationships between the level of circulating fibrocytes and the density of CD45^+^ FSP1^+^ cells in distal airways measured in control subjects (black circles) and COPD patients (open circles). B, C, Correlation coefficient (r) and significance level (P value) were obtained by using nonparametric Spearman analysis.

**Fig E10.**
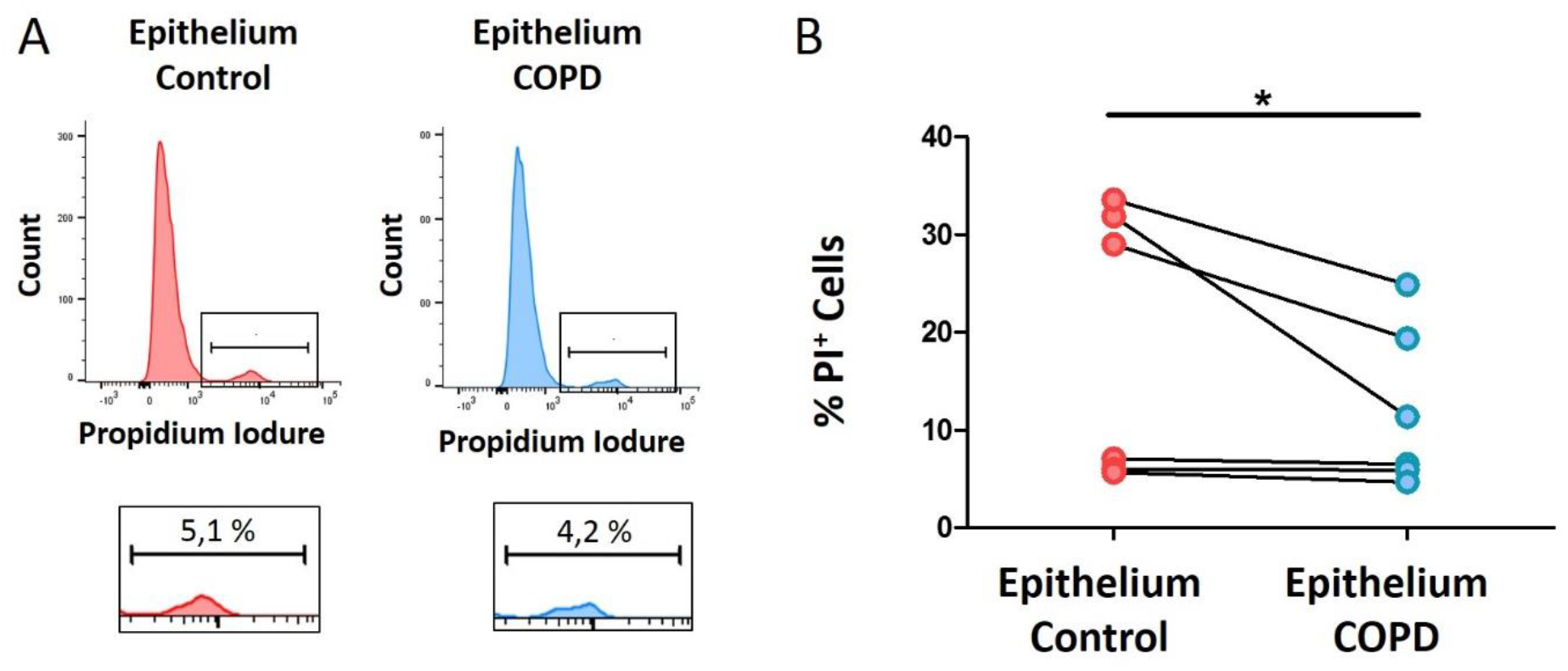
Influence of epithelium on fibrocyte survival. A, Representative histograms of flow cytometry for Propidium Iodure (PI) fluorescence recorded on fibrocytes exposed to epithelium supernatants, either from control subjects (left panel), or COPD patient (right panel). A gate was drawn to define the population of dead cells (PI^+^ cells). B, Quantification of PI^+^ cells in fibrocytes from COPD patients (n=6) exposed to epithelium supernatants from control subjects (red circles) or COPD patients (blue circles). *: P<0.05, Wilcoxon test.

## Abbreviations

APC: Allophycocyanin
BEC: Bronchial Epithelial Cells
CSA: Cross Section Area
CSN: Cross Section Number
GOLD: Global Initiative for Chronic Obstructive Lung Disease
COPD: Chronic Obstructive Pulmonary Disease
CT: Computed tomography
FEV_1_: Forced Expiratory Volume in 1 second
FITC: Fluorescein isothiocyanate
FVC: Forced Vital Capacity
FSP1: Fibroblast-Specific Protein 1
LA: Lumen Area
LAA: Low Attenuation Area
MLA: Mean Lung Attenuation
PaCO_2_: Arterial Partial Pressure of Carbon Dioxide
PaO_2_: Arterial Partial Pressure of Oxygen
PE: Phycoerythrin
PBMC: Peripheral Blood Mononuclear Cells
TLCO: Transfer Lung capacity of Carbon monoxide
WA: Wall Area
WT: Wall Thickness

